# Structure-guided engineering of CCL27 enhances natural ligand CAR T-cells against CCR10 for multiple myeloma

**DOI:** 10.1101/2025.08.19.670119

**Authors:** Nikhil Chilakapati, Bonell Patiño-Escobar, Emily Y. Chen, Radhika Dalal, Jonathan de Montagnac, Haley Johnson, Amrik S. Kang, Naomi Akagi, Emilio Ramos, Paul Phojanakong, Fernando Salangsang, Yan Zeng, Ajai Chari, Alfred Chung, Anupama D. Kumar, Thomas G. Martin, Jeffrey L. Wolf, Brian R. Shy, Veronica Steri, William J. Karlon, Tanja Kortemme, Benjamin G. Barwick, Arun P. Wiita

## Abstract

Despite the success of BCMA CAR-Ts, many multiple myeloma patients relapse and require additional therapeutic options. Our group previously identified the chemokine receptor CCR10 as a potential alternate target to address this need. Here, we validated CCR10 expression on primary myeloma tumors and sought to develop CAR T-cells against CCR10, utilizing its natural ligand CCL27 as a CAR binding element. However, CARs based on the native CCL27 sequence were ineffective. We thus utilized computational modeling and structure-guided engineering to inform rational mutations along the CCL27-CCR10 interface, exploiting a hydrophobic pocket on CCR10. This effort identified CCL27 mutants with an additional N-terminal aromatic amino acid that dramatically improved the efficacy of CCL27-based CAR-Ts to near that of current anti-BCMA CAR-Ts. We validated key amino acid contacts at the CCL27-CCR10 interface, which contribute to increased CAR binding avidity, predicted to be influenced by increased Van der Waals interactions. Lastly, we found that the CCL27 mutants have no toxicity in the hematopoietic compartment. This work illustrates the potential of engineering natural ligand CAR-Ts beyond their wild-type sequences and underscores the translational potential of engineered CCL27 mutant CAR-Ts.

## Introduction

Multiple myeloma (MM) is a plasma cell malignancy that remains without a long-term cure despite recent therapeutic advances^1^. One of the most promising of these novel treatment options are chimeric antigen receptor (CAR) T-cells targeting the surface protein B-cell maturation antigen (BCMA)^2^. Despite the FDA approval of two highly effective anti-BCMA CAR-Ts, idecabtagene vicleucel (“ide-cel”) and ciltacabtagene-autoleucel (“cilta-cel”), the vast majority of patients eventually relapse^1,3^. Although not observed ubiquitously, relapsed tumors of patients after anti-BCMA therapies often show a loss of surface BCMA expression^3,4^. The prognosis of salvage regimens for these patients who relapse after current anti-BCMA CAR-Ts is quite poor^5,6^. Additionally, anti-BCMA CAR-Ts have seen neurotoxicity that takes the form of delayed Parkinsonism in up to 5% of patients treated with the most widely adopted and efficacious CAR-T product, cilta-cel, hypothesized to be caused by the potential expression of BCMA in the basal ganglia^7,8^. There have also been recent reports of acute gut toxicities from BCMA CAR-Ts, termed immune effector cell-associated enterocolitis, at around 2% incidence for cilta-cel^9^. Thus, there is a significant unmet need for therapeutic targets for multiple myeloma beyond BCMA, which ideally have fewer toxicities.

To address this issue, many groups, including ours, have set out to identify additional CAR-T targets for multiple myeloma. The most advanced of these novel targets is the orphan G protein-coupled receptor (GPCR), G protein-coupled receptor, class C, group 5, member D (GPRC5D), which has shown promising results in preclinical studies and a phase 1 clinical trial^10,11^. However, responses do not yet appear durable, with a median duration of response of 7.8 months^11^. Additionally, GPRC5D also has expression in a subset of cells in the hair follicles of the skin, hard keratinizing tissue, and potential expression in the cerebellum or inferior olivary nucleus, with anti-GPRC5D CAR-Ts showing toxicities in these tissues, as well as significant dysgeusia for many patients^11^. Therefore, there remains a need for additional therapeutic targets for multiple myeloma that have minimal off-target expression with products that have long-lasting responses.

To this end, our group previously performed comprehensive surface proteomics of myeloma cell lines and primary samples, creating a ranking of surface antigens for their potential as therapeutic targets^12^. We specifically focused on targets with high expression on plasma cells but minimal expression on any other normal tissues.

Filtering through the hits of this analysis, we focused on CCR10 as it scored very high in our ranking but had yet to be explored as a therapeutic target for multiple myeloma^12^.

Additionally, via analysis of the Multiple Myeloma Research Foundation (MMRF) CoMMpass database^13^, we found that *CCR10* was upregulated in high-risk disease and associated with poorer clinical outcomes^12^. CCR10 is a chemokine receptor with expression primarily on subsets of lymphocytes, particularly plasma cells and skin-homing T-cells, where it plays a role in lymphocyte migration to specific tissues^14,15^.

CCR10 interacts with its ligands CCL27 and CCL28 to perform its functions in cell motility^16^. Like the other chemokine receptors, CCR10 is a GPCR and a seven-pass transmembrane protein^15^. The complex membrane-embedded topology of GPCRs is well-known to create significant challenges in antibody discovery pipelines^17^.

Chemokine receptors often have high sequence homology because they interact with many of the same chemokines, leading to difficulties in generating highly selective antibody reagents^18,19^. To circumvent these issues, we previously attempted to use CCR10’s natural ligand, CCL27, as the binder for a CAR construct^12^; however, we found that these CAR-Ts had very limited anti-tumor efficacy and were not appropriate for further therapeutic development. We specifically chose CCL27 to avoid off-target effects, as the other ligand, CCL28, is known to also interact with the more widely expressed receptor CCR3^20^.

Notably, several other natural ligand-based CAR T-cells have been described, pursuing targets ranging from BCMA in myeloma^21^ to CD70 in Acute Myeloid Leukemia^22^ and myeloma^23^ to CD4 in HIV^24^. Current natural ligand designs based on wild-type (WT) ligand sequences often do not appear able to achieve the same efficacy as CARs based on antibody fragments (scFv’s or nanobodies)^25–27^, which enable exploration of a much broader sequence and affinity space^28–30^. Yet natural ligand CAR designs are attractive to consider as they are likely to be less immunogenic than antibody fragments and may be able to interact with physiologically relevant epitopes not accessible by antibody fragments. However, a major unanswered question in the field is whether protein engineering strategies can be used to improve the activity of these natural ligand CARs beyond evolutionarily defined constraints.

Here, we sought to address this challenge by applying a structure-guided design approach to CCL27-based, anti-CCR10 CAR-Ts. We identified two N-terminal extensions of CCL27 that significantly increased the efficacy of our CAR-Ts to near the level of the current clinically approved anti-BCMA CAR-Ts. We also identified a potential mechanism for the increased functionality of the CCL27 mutant CAR-Ts *in silico*, with validation by functional studies. Lastly, we performed comprehensive toxicity testing of our CCL27 mutant CAR-Ts in the peripheral blood compartment, finding no evidence of off-tumor toxicity. Collectively, our work identifies a promising new therapeutic for multiple myeloma while also demonstrating that structure-guided engineering approaches can be used to enhance natural-ligand CAR designs.

## Results

### CCR10 is an intriguing therapeutic target for multiple myeloma

Building off our initial work identifying CCR10 as a new immunotherapy target for myeloma, we sought further validation on the therapeutic potential of targeting CCR10. In our prior work, we showed ubiquitous CCR10 expression on the CD138+ tumor cell fraction of 10 myeloma primary samples^12^. Here, we performed flow cytometry on 8 additional primary myeloma samples and again found consistent CCR10 expression (**Fig. 1A**). We further characterized several common myeloma cell lines and identified that they are all positive for CCR10 (**Fig. 1B**). We also re-analyzed a published single-cell RNA sequencing (scRNA-seq) dataset^31^ comparing myeloma vs healthy donor bone marrows, confirming that *CCR10* is upregulated in plasma cells from myeloma patients and that *CCR10* expression is restricted to the plasma cell cluster, similar to BCMA (*TNFRSF17*) (**Fig. 1C**). This result was validated in an independent scRNA-seq cohort^32^, where *CCR10* expression is significantly upregulated in myeloma plasma cells as compared to healthy donors (**Supplementary Data Fig. 1A**). scRNA-seq data from normal tissues compiled in the Human Protein Atlas^33^ show that *CCR10* is strongly expressed in plasma cells, with minimal expression in other cell types (**Fig. 1D**). We also analyzed the recently described subtypes of myeloma^13,34^ in the MMRF CoMMpass dataset for *CCR10* expression and found that the highest risk PR subtype has strong expression of *CCR10* as compared to the other lower risk subtypes (**Fig. 1E**).

**Figure 1:**
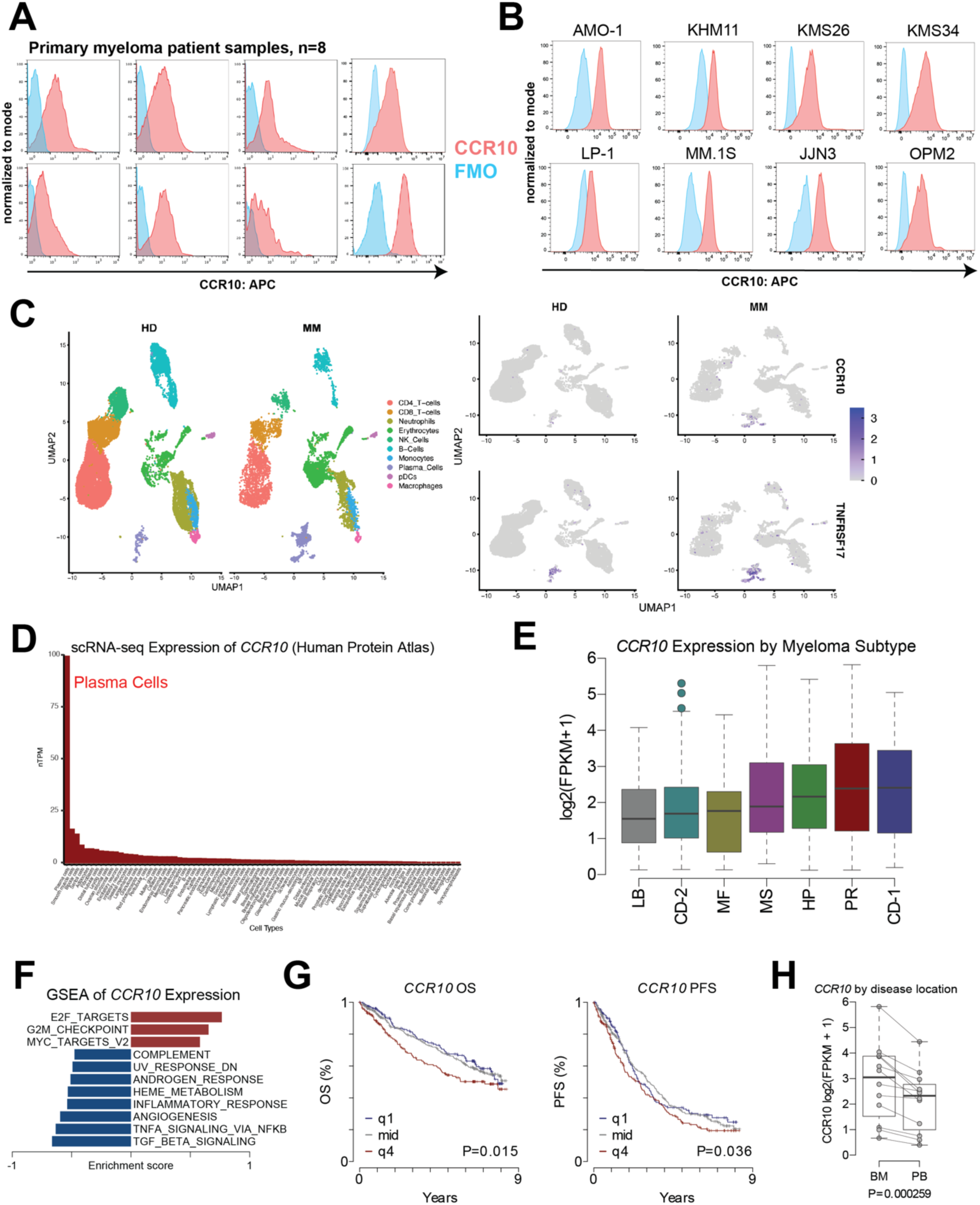
CCR10 expression is primarily restricted to myeloma plasma cells with upregulation in high-risk disease. **A.** Flow cytometry profiling of CCR10 on fresh MM patient tumor samples, gating on myeloma cells (CD19-& CD138+ or CD38+ & CD45 dim/-& CD19-) as compared to FMO (Fluorescence Minus One) negative control (*n* = 8). **B.** Flow cytometry on a panel of 8 MM cell lines shows uniform CCR10 positivity. **C.** Single-cell RNA sequencing of 9 myeloma bone marrows with >25% of plasma cells compared with 7 healthy donor bone marrows (GSE223060). **D.** Single-cell RNA sequencing across human cell types taken from the Human Protein Atlas^33^. **E.** Expression of *CCR10* in previously annotated gene expression subtypes in newly diagnosed multiple myeloma: (LB: Low bone disease; CD-2: Cyclin D1 and CD20; MF: MAF; MS: MMSET; HP: Hyperdiploid; PR: Proliferation; CD-1: Cyclin D1; *n* = 764). **F.** Gene Set Enrichment Analysis (GSEA) of Hallmark gene sets correlated with *CCR10* expression with an FDR < 0.01. **G.** Progression-Free Survival (PFS; left) and Overall Survival (OS; right) stratified by *CCR10* expression quartiles (q1 low: blue; mid quartiles: gray; q4 high quartile: red). *p*-values were calculated using Cox Proportional hazards regression Wald test, treating *CCR10* expression as a continuous variable (*n* = 764). **H.** *CCR10* expression in paired bone marrow (BM) and Peripheral Blood (PB) CD138+ MM cells (*n* = 24 samples; 11 patients). *p*-values were calculated using linear regression with a covariate for patient. Analyses were performed on data from the MMRF CoMMpass dataset^13^ (**E-H**).

Additionally, we found that *CCR10* expression in myeloma was correlated with gene programs associated with cell cycle and proliferation (**Fig. 1F**). We further identified that the top quartile (q4) of *CCR10* expressors had significantly worse survival outcomes, by both overall survival (OS) and progression-free survival (PFS) (**Fig. 1G**). *CCR10* expression was also found to be an independent risk factor in myeloma, with significant upregulation in relapsed/refractory (RR) as compared to patient-matched newly diagnosed (ND) tumors (**Supplementary Data Fig. 1B-C**). Interestingly, we found that *CCR10* expression was significantly lower in myeloma plasma cells isolated from peripheral blood as compared to the bone marrow of the same patients, indicating a potential role of CCR10, as a chemokine receptor, in the homing and maintenance of myeloma plasma cells in the bone marrow (**Fig. 1H**). Together, these findings further highlight CCR10 as a promising immunotherapy target for myeloma and high-risk patients with aggressive disease in particular.

### Structure-guided mutations to CCL27 identify promising CAR design candidates

Consistent with our prior results, we first confirmed that WT CCL27-based CAR-Ts had minimal *in vitro* anti-myeloma efficacy at lower effector:tumor (E:T) ratios (**Supplementary Data Fig. 2A**). This finding motivated us to undertake further engineering of this binder to create a clinically translatable therapeutic. We hypothesized that the ineffectiveness of the CCL27-based CAR-T may be due to low binding affinity and/or avidity. Prior studies of an anti-BCMA CAR-T based on the natural ligand APRIL^21^ suggested that designing a trimeric ligand binder could overcome this hurdle via improved avidity (**Supplementary Data Fig. 2B**). However, this approach yielded little to no improvements in the context of our CCL27-based CAR-Ts (**Supplementary Data Fig. 2C**). Refocusing our efforts, we combined insight from AlphaFold3 predicted structural models with literature descriptions of the chemokine-chemokine receptor binding interface. AlphaFold3 was utilized because it was trained on structurally similar solved chemokine-chemokine receptor structures, generating relatively high-confidence CCL27-CCR10 predicted structures^35^. The literature on the two-step model^36^ of chemokines binding their receptors enabled us to annotate and validate the key binding interfaces of the AlphaFold3 structural predictions (**Fig. 2A-B**). These insights motivated us to design a small library of mutations to CCL27 at the highlighted functional domains (**Fig. 2B-C**). We particularly focused on the N-terminus of CCL27 and identified potential unoccupied residues at the CS2-CRS2 interface (**Fig. 2A**). Notably, by rationally testing several N-terminal extensions of the WT sequence based on our structural models, we explored a structural and sequence space that would not be accessible to a traditional affinity maturation campaign for a target binder^37^. Screening this library for *in vitro* cytotoxicity, we identified two mutations that add a single N-terminal aromatic amino acid, tryptophan (W-CCL27) or phenylalanine (F-CCL27), that remarkably increased the efficacy of CCL27-based CAR-Ts from essentially equal to used negative controls to near the level of the positive control anti-BCMA CAR-T (cilta-cel binder), in one or both myeloma cell lines tested (**Fig. 2D**).

**Figure 2:**
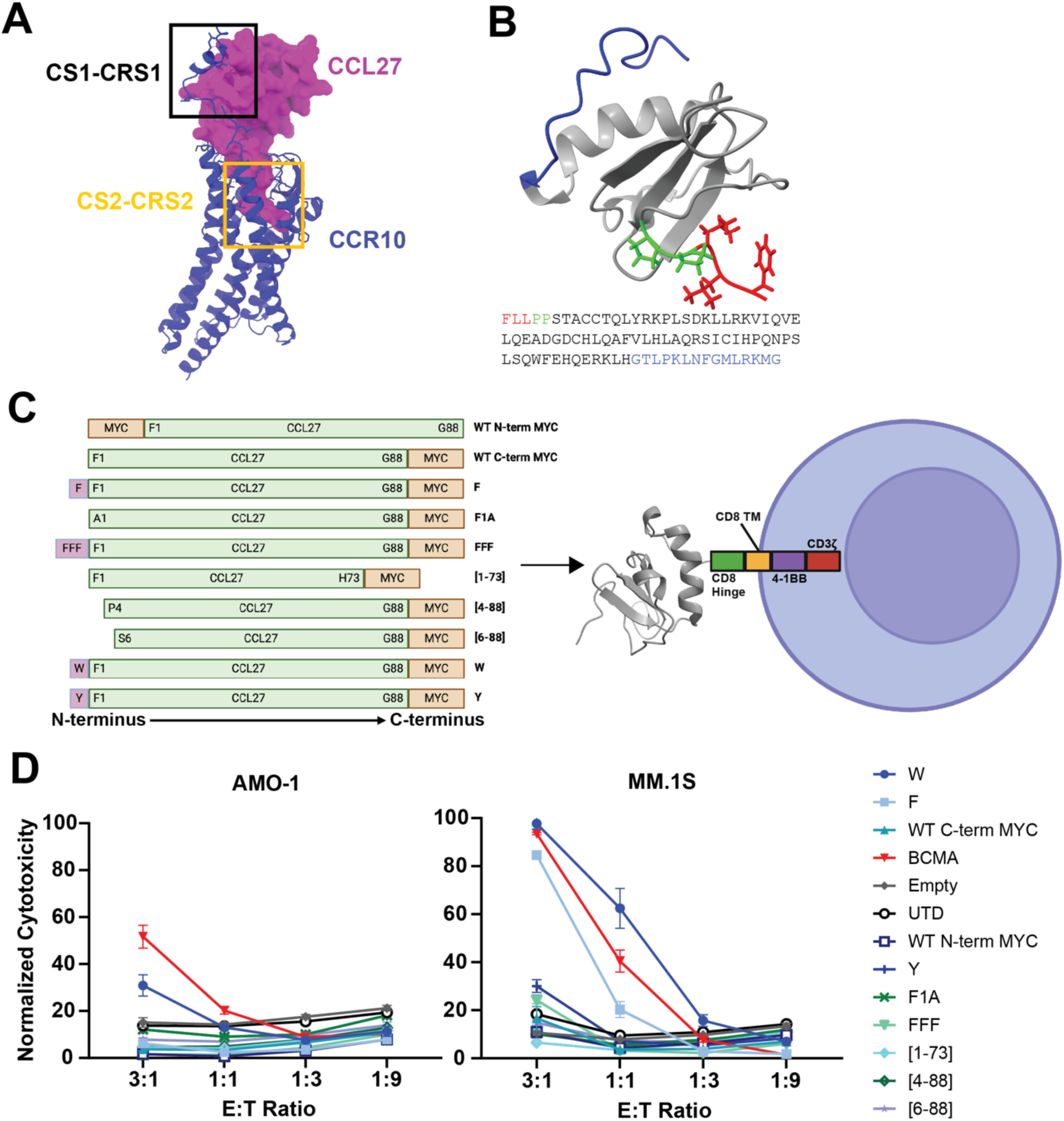
Structure-guided investigation of CCL27-CCR10 binding identifies key interfaces and informs CCL27 mutations. **A.** CCL27-CCR10 binding interaction modeled in AlphaFold3 and displayed using ChimeraX^53^, with two key binding interfaces highlighted (CS1-CRS1 & CS2-CRS2). **B.** Key binding interfaces of CCL27 from AlphaFold3 predicted models highlighted on the CCL27 NMR structure and amino acid sequence. **C.** Illustration of CCL27 mutational library implemented as the binder into a standard CAR-T backbone. **D.** *In vitro* cytotoxicity assays comparing tumor lysis of various CCL27 mutants with positive control of anti-BCMA CAR and negative controls of Empty CAR and UTD (untransduced T-cells) at the indicated effector:tumor (E:T) ratios (*n* = 3 technical replicates). After 18 hours of co-culture, cytotoxicity was measured by incubation with luciferase and readout by luminescence.

These surprising results indicated that we could achieve significant CAR-T enhancement by rationally adding just a single amino acid to the N-terminus of CCL27. Next, we confirmed the robust efficacy of the W-CCL27 and F-CCL27 mutants across additional T-cell donors and three different CCR10+ myeloma cell lines (MM.1S, JJN-3, AMO-1), which was yet again near the level of the anti-BCMA CAR-T positive control (**Fig. 3A**). Additionally, no cytotoxicity was seen in the *CCR10* knockout AMO-1 myeloma cell line, supporting that our mutants did not alter the specificity of CCL27 for CCR10 (**Fig. 3A**). In line with this result, we also noted a somewhat decreased efficacy of our mutant CCL27 CAR T-cells against additional myeloma lines with low CCR10 expression (**Supplementary Data Fig. 2D**). Furthermore, we found that both CD4+ and CD8+ mutant CCL27 CAR T-cells proliferate equally to anti-BCMA CAR-Ts in response to MM.1S co-culture (**Fig. 3B & Supplementary Data Fig. 3A**). We also found that after tumor exposure, mutant CCL27 CAR-Ts were distinctly less exhausted than the anti-BCMA CAR-Ts, accentuated by a more central memory phenotype and a lower expression of the exhaustion markers LAG-3, PD-1, and TIM-3 (**Fig. 3C & Supplementary Data Fig. 3B-C**). Additionally, we observed that the mutant CCL27 CAR-Ts had significantly lower secretion of traditional effector cytokines (GM-CSF, IFNg, and TNFa) than the anti-BCMA CAR-T, yet significantly higher than the WT-CCL27 and negative control empty CAR (**Supplementary Data Fig. 3D**). Similar trends were present for Th2-type cytokines (IL-4, IL-5, IL-6, IL-10, IL-13) (**Supplementary Data Fig. 3D**). Notably, across these T-cell phenotype characterization assays, we also observed that the WT-CCL27 CAR-T, while showing no meaningful anti-tumor cytotoxicity *in vitro*, did display some signatures of activity (**Fig. 3B-C & Supplementary Data Fig. 3D**). These findings motivated us to characterize the ability of the mutant CCL27 CAR-Ts to handle repetitive antigen stimulation. Results showed that the mutant CCL27 and anti-BCMA CAR-Ts were able to control the MM.1S tumor after three repetitive stimulations, while the WT-CCL27 CAR-Ts rapidly lost control of the tumor (**Fig. 3D**). In terms of CAR T-cell proliferation, the mutant CCL27 CAR-Ts had robust initial proliferation that was reduced upon subsequent stimulation, while the anti-BCMA CAR-Ts showed proliferation at all stimulations (**Supplementary Data Fig. 3E**). Lastly, we confirmed the ability of our CAR-Ts to effectively kill primary CCR10+ myeloma cells *ex vivo* from two independent patients (**Fig. 3E & Supplementary Data Fig. 3F**). Taken together, these results show that the rational addition of just a single aromatic amino acid at the N-terminus of CCL27 can significantly improve the efficacy of CCL27-based CAR-Ts to near the level of the cilta-cel anti-BCMA CAR-T in numerous CAR-T characterization assays.

**Figure 3:**
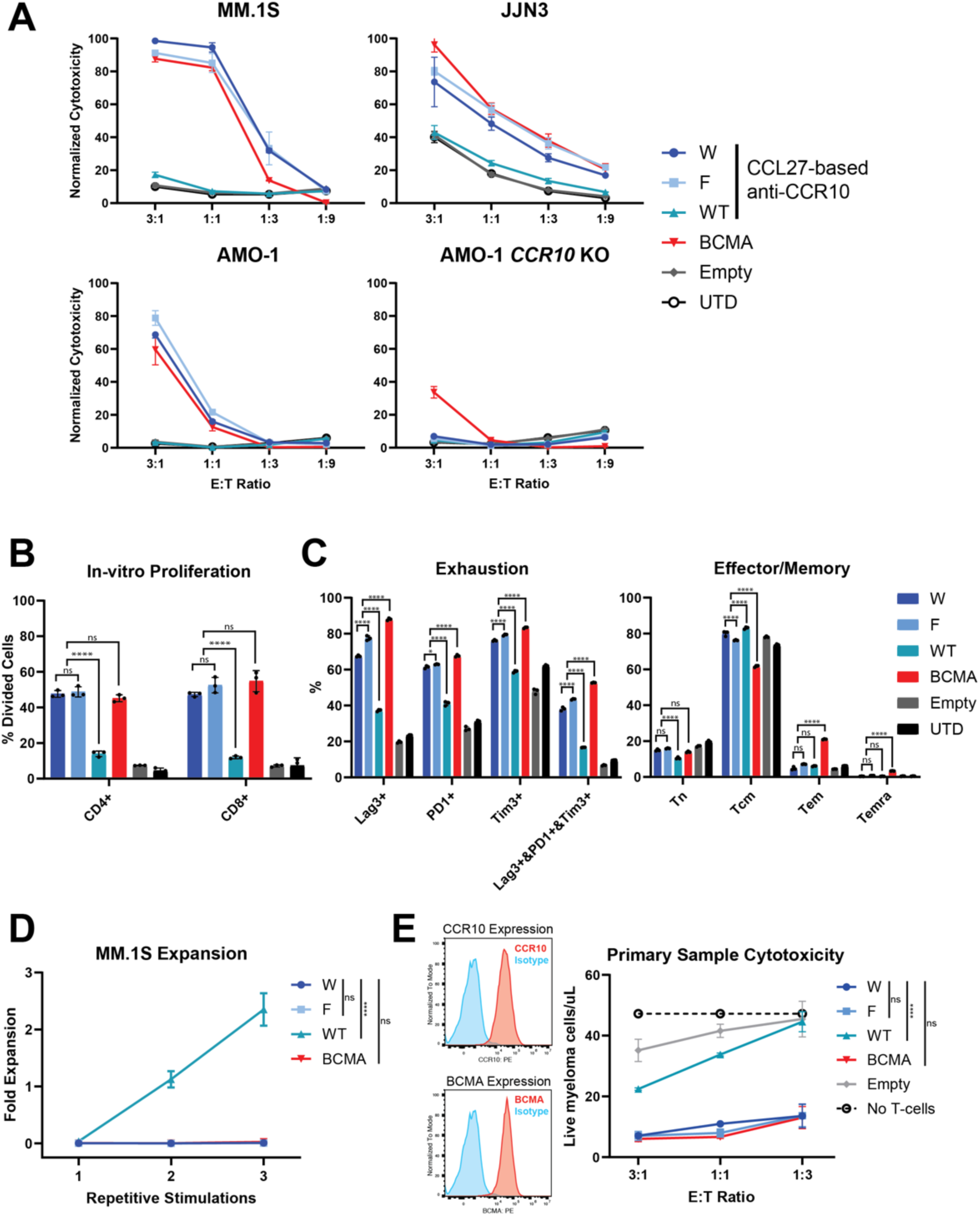
Characterization of mutant CCL27 CAR T-cells. **A.** *In vitro* cytotoxicity after 18 hours of co-culture of the previously identified mutant CCL27 CAR-Ts with CCR10+ MM cell lines (MM.1S, JJN3, AMO-1) and the CCR10-negative AMO-1 *CCR10* KO cell line. The anti-BCMA CAR was a positive control, and Empty CAR and UTD (untransduced T-cells) were negative controls at the indicated effector:tumor (E:T) ratios (*n* = 3 technical replicates). Data presented is representative of *n = 2* T-cell donors, but only one T-cell donor is shown. **B.** *In vitro* proliferation after 72 hours of co-culture with the MM.1S tumor cells as measured by labeling CAR-Ts with Cell Trace Violet and gating “Divided Cells” as cells that have undergone at least one division. *p*-values calculated by 2-way ANOVA with Tukey’s correction for multiple comparisons applied: ns = not significant & **** = *p*<0.0001. **C.** T-cell phenotype after 24-hour co-culture with MM.1S tumor cells measured by flow cytometry, with *n* = 3 technical replicates, characterizing exhaustion by percent of cells expressing the exhaustion markers of Lag-3, PD-1, and Tim-3. Effector-memory state was calculated by four populations of expression of markers CD62L and CD45RA. *p*-values calculated by 2-way ANOVA with Tukey’s correction for multiple comparisons applied: ns = not significant, * = *p*<0.05, & **** = *p*<0.0001. **D.** Repetitive stimulation of CAR-Ts with tracking of tumor cell fold expansion between stimulations by flow cytometry after 48 hours of co-culture. *p*-values calculated by 2-way ANOVA: ns = not significant & **** = *p*<0.0001. **E.** Primary sample *in vitro* cytotoxicity assay using flow cytometry to identify the depletion of myeloma cells (CD19-& CD138+), with count beads utilized for absolute quantification of *n* = 3 technical replicates and representative of *n* = 2 primary samples and T-cell donors. *p*-values calculated by 2-way ANOVA: ns = not significant & **** = p<0.0001.

### CCL27 mutant CAR-Ts are highly functional in myeloma xenograft models

After characterizing the highly promising *in vitro* performance of the CCL27 mutant CAR T-cells, we moved forward to profile their functionality *in vivo*. In an initial pilot study, we first chose the MM.1S myeloma xenograft model with three NSG mice per arm. We found that both CCL27 mutant CAR-Ts showed improved tumor control over WT-CCL27 and Empty CAR-Ts, similar to the BCMA CAR-T, consistent with our *in vitro* findings (**Fig. 4B-C**). In this small-scale study, we were also encouraged to find a trend toward increased survival (*p*=0.06 by log-rank test) by the CCL27 mutant CAR T-cells compared to WT-CCL27 and Empty CAR (**Fig. 4C-D**).

**Figure 4:**
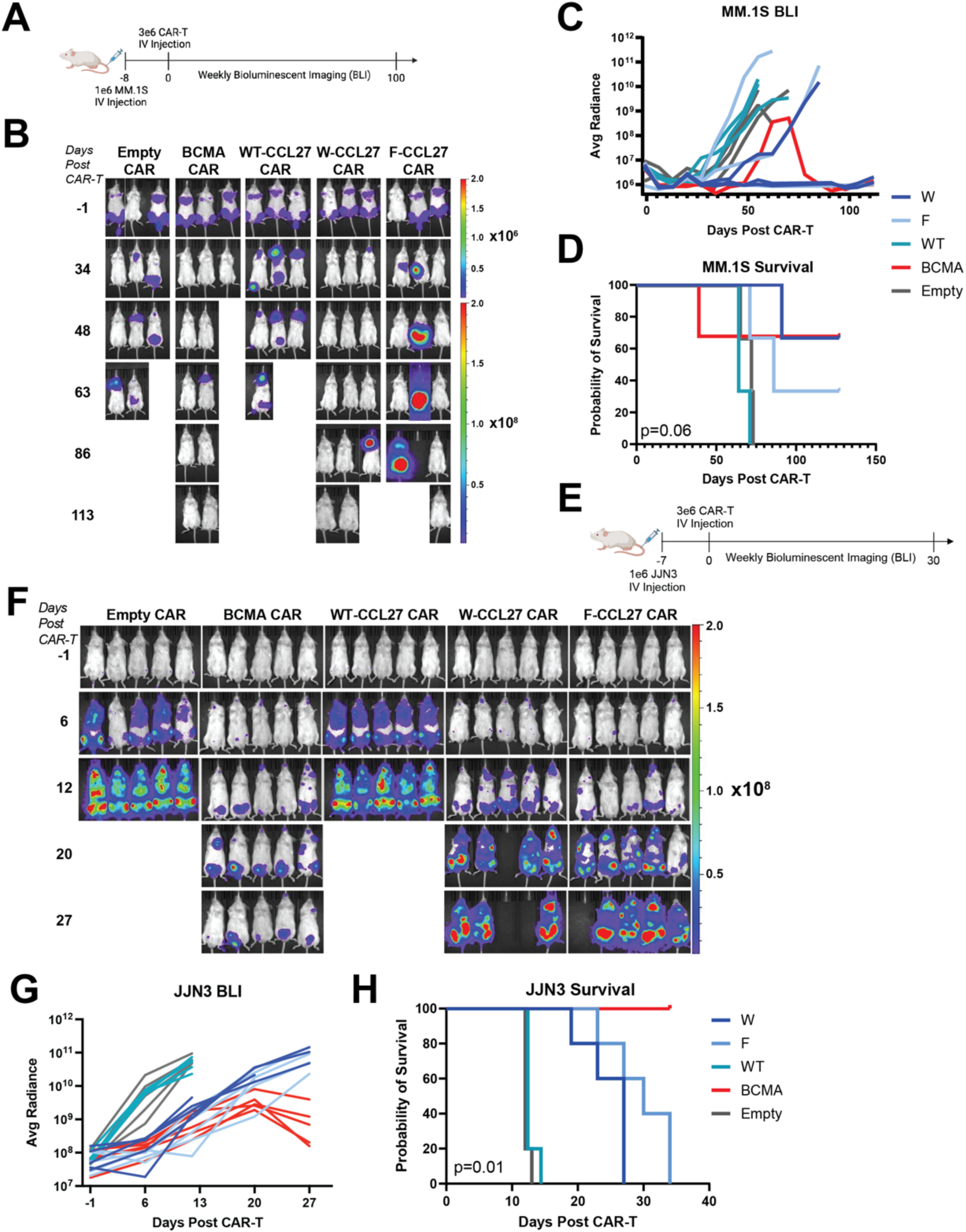
I*n vivo* functionality of mutant CCL27-based CAR-Ts. **A.** Illustration of MM.1S murine study design with MM.1S cells orthotopically implanted in NSG mice, *n* = 3/arm. **B.** Weekly bioluminescent imaging of CCL27-based CAR-Ts (W-, F-, and WT-CCL27) compared with positive control BCMA CAR and negative control Empty CAR. Note that one mouse in the BCMA CAR arm was required to be sacrificed without any evidence of tumor burden, likely due to graft-versus-host disease (GvHD) based on symptoms. **C.** Quantified bioluminescence imaging data of the MM.1S study illustrating tumor control for CCL27 mutant CAR-Ts, as compared to WT-CCL27 and Empty CARs. Certain datapoints were excluded when clear spillover of bioluminescence between mice was observed. **D.** Kaplan Meier curve of survival from the MM.1S murine study as tested by log-rank test (Mantel-Cox). **E.** Illustration of JJN3 murine study design with JJN3 cells orthotopically implanted in NSG mice, *n* = 5/arm. **F.** Weekly bioluminescent imaging of the aggressive JJN3 tumor model with the same arms as the previous MM.1S study. **G.** Quantified bioluminescent imaging of the JJN3 study showing a slowed tumor progression provided by the CCL27 mutant CAR-Ts as compared to the WT-CCL27 and Empty CAR-T. **H.** Kaplan Meier curve of the survival of mice in the JJN3 murine study was tested by the log-rank test (Mantel-Cox).

Following these pilot results in the relatively indolent MM.1S model, we moved forward to perform a stress test experiment in the more aggressive JJN3 myeloma model. After 7 days, a very high tumor burden (bioluminescence radiance between 10^7^-10^8^ photons/s/cm^2^/sr) was observed prior to IV implantation of 3e6 CAR+ T-cells per mouse (**Fig. 4E**). Under this aggressive model, we observed rapid tumor growth, where the humane survival endpoint was reached for both the Empty CAR negative control and WT-CCL27 CAR in under two weeks (**Fig. 4F-H**). CCL27 mutant CARs initially performed equivalently to the BCMA CARs in terms of tumor control (**Fig. 4F-G**), though the BCMA CAR response was ultimately more durable. However, we remained encouraged that our mutant CAR Ts significantly improved survival vs. WT-CCL27 CAR in this highly aggressive tumor model (**Fig. 4H**) (*p*=0.01 by log-rank test). These *in vivo* results further supported the dramatic potency enhancement driven by a single N-terminal amino acid addition to CCL27.

### Mechanism of CCL27 mutants binding explains enhanced anti-CCR10 CAR-T efficacy

We next explored the mechanism of why the addition of a nonpolar aromatic amino acid (W or F) at the N-terminus of CCL27 significantly improved the efficacy of CCL27-based CAR-Ts. To help answer this, we performed an in-depth characterization of the AlphaFold3 structural models to identify a binding pocket, consisting of four critical amino acids (V47, S48, Y120, L300), on CCR10 that are uniquely accessed by the CCL27 mutants (**Fig. 5A**). Aiming to functionally validate this model prediction, we mutated these four amino acids to either alanine (A), lysine (K), or tryptophan (W) to selectively disrupt the interactions at the binding interface of the CCL27 mutants to varied extents. These mutant forms of CCR10 were then expressed in AMO-1 cells with *CCR10* KO and confirmed by flow cytometry that all CCR10 mutants, except V47K, were stably expressed with a similar antigen density (**Supplementary Data Fig. 4A**).

**Figure 5:**
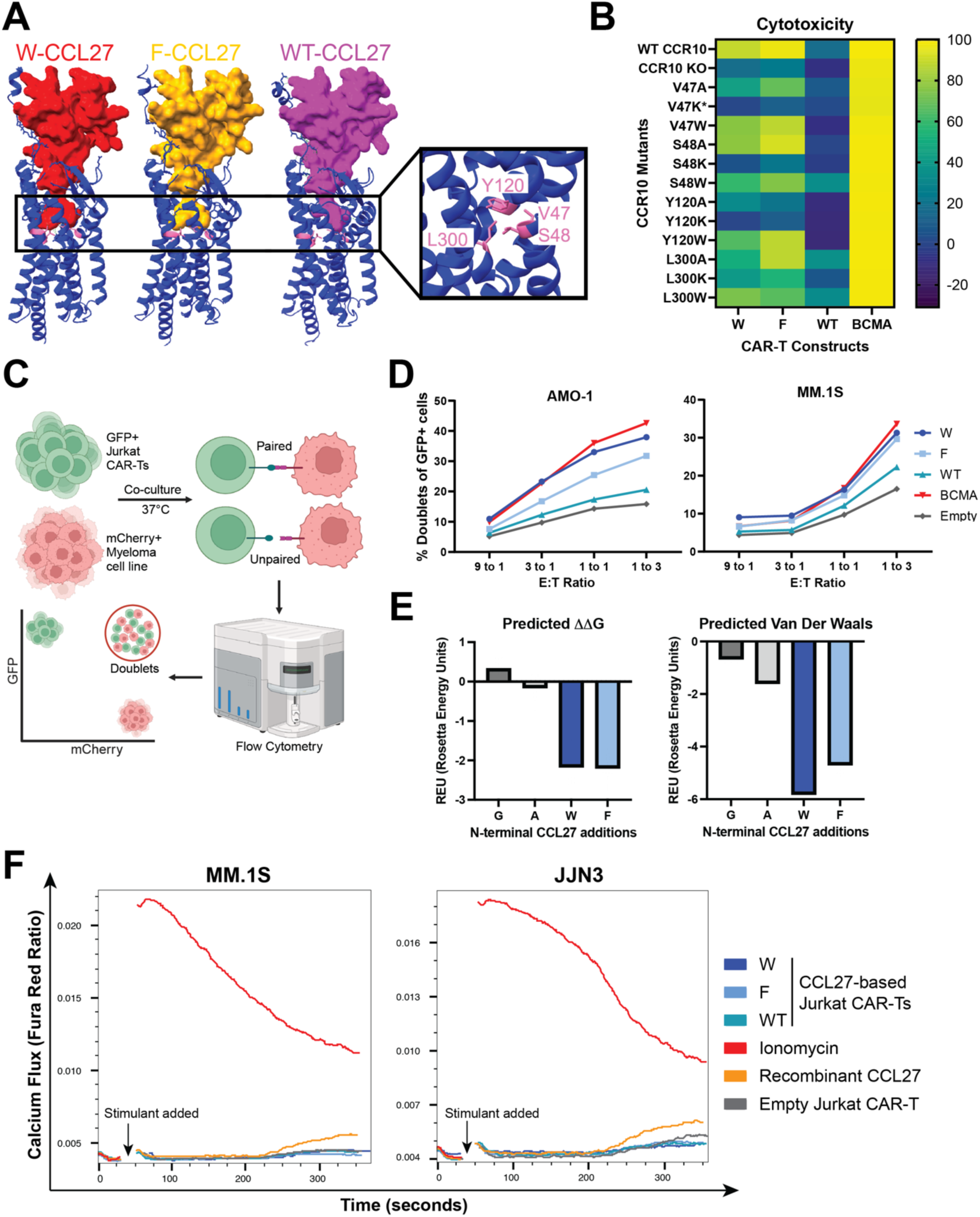
Investigating the mechanism of CCL27 mutants binding CCR10. **A.** AlphaFold3 models of binding between W-CCL27 (red), F-CCL27 (yellow), and WT-CCL27 (purple) binding CCR10 (blue). The binding pocket of the N-terminal residues of CCL27 is highlighted in the black box, with the amino acids found on CCR10 to uniquely contact the CCL27 mutants colored in pink, with their identity shown. **B.** Cytotoxicity of the CAR-Ts on the x-axis (W-CCL27, F-CCL27, WT-CCL27, BCMA) against AMO-1 cell line engineered to express the forms of CCR10 (WT, KO, or mutants) on the y-axis at an 8:1 E:T ratio. Cytotoxicity values are representative of *n* = 3 technical replicates and normalized by the cytotoxicity of UTD T-cells against their respective target cell line. Clear outlier values were excluded for graphical representation. **C.** Illustration of the doublet assay protocol used for assessing the avidity of Jurkat CAR-Ts binding myeloma tumor cells. **D.** Percent of doublets (mCherry+ & GFP+), representative of *n* = 2 technical replicates, gated on the GFP+ Jurkat CAR-Ts for the AMO-1 and MM.1S myeloma cell lines at the shown E:T ratios. **E.** Relevant metrics outputted from the Rosetta FlexddG protocol using the fa_talaris2014 scoring function on the AlphaFold3 structural predictions of N-terminal extensions to CCL27 binding CCR10. (*Left panel*) Predicted ΔΔG values outputted in Rosetta Energy Units (REU) as compared to the G-CCL27 binding CCR10 structure. (*Right Panel*) Predicted energy contributions due to van der Waals interactions as compared to the G-CCL27 binding CCR10 structure. Each bar is representative of *n* = 20 output structures generated by FlexddG. **F.** Ratiometric analysis of the Fura Red calcium flux dye^54^ signal of labeled myeloma cell lines (MM.1S and JJN3) off the blue laser (Ca^2+^ free) divided by the violet laser (Ca^2+^ bound). Values shown depict the moving average of the Fura Red ratio over time (∼5 min) of a single replicate but are representative of three independent experiments. The arrow indicates the time that the stimulants in the legend were added.

Cytotoxicity of our CCL27 mutant CAR T-cells against these mutant CCR10 cell lines showed that mutations at the V47, S48, and Y120 amino acids on CCR10 uniquely lowered the efficacy of the CCL27 mutant CAR-Ts, particularly the lysine mutations, indicating the importance of these amino acids for the functionality of the CCL27-based CAR-Ts (**Fig. 5B**). We next adapted a recently described flow cytometry-based avidity assay^38^ to assess the binding avidity of the CCL27 mutant CAR-Ts in an on-cell manner, which involved co-culturing GFP+ Jurkat T-cells expressing the CAR with mCherry+ myeloma cells and reading out cell-cell interactions as GFP+ & mCherry+ doublets (**Fig. 5C & Supplementary Data Fig. 4B-C**). We note that Jurkat cells are used instead of primary T-cells here as Jurkat T-cells do not have cytotoxic capacity when expressing a CAR and, therefore, can generate stable doublets instead of eliminating tumor cells.

The assay showed that the CCL27 mutants bound myeloma cells more strongly than WT-CCL27 in both the MM.1S and AMO-1 cell lines, and nearly equal to that of the cilta-cel anti-BCMA binder (**Fig. 5D**). Further computational analysis of the AlphaFold3 predicted structures with the Rosetta-based Flex ddG protocol^39^ showed that the addition of a nonpolar aromatic residue (W or F) to CCL27 stabilized the CCL27-CCR10 interface and showed more van der Waals interactions compared to the addition of a minimal glycine or alanine residue (**Fig. 5E**).

Given that soluble CCL27 can stimulate downstream signaling via CCR10 binding^15,40^, and the increased expression of CCR10 in highly proliferative myeloma tumors and its association with proliferative gene programs (**Fig. 1E-G**), we next became curious whether the binding of CCL27 mutant CARs could influence possible CCR10-based signaling and growth in myeloma cells. First, we found that the CCL27 mutants, expressed as Jurkat CAR-Ts, were not agonistic of CCR10 signaling, by calcium flux assays in co-culture with myeloma cell lines, as compared to the recombinant CCL27 and Ionomycin controls, which showed appreciable calcium flux signaling (**Fig. 5F**). Exploring the downstream effects of CCR10 signaling, we found that co-culturing the CCL27 mutant Jurkat CAR-Ts with myeloma cell lines had minimal effects on two canonical signaling effectors of chemokine receptors, p-Akt and p-Erk from the PI3K and MAPK signaling pathways respectively (**Supplementary Data Fig. 4E**). Lastly, co-culturing Jurkat CCL27 mutant CAR-Ts with various myeloma cell lines did not meaningfully alter myeloma growth across 72 hours (**Supplementary Data Fig. 4F**). Together, these findings are consistent with our initial hypothesis that these N-terminal extensions of CCL27 improve the energetics of CCR10 binding, particularly by van der Waals interactions, ultimately driving improved CAR-T efficacy. However, we do not anticipate that interactions of our engineered CCL27-based CAR-Ts with CCR10+ myeloma will stimulate meaningful CCR10 signaling or tumor growth.

### CCL27 mutant CAR T-cells have no toxicity in the peripheral blood

To ascertain the therapeutic potential of our anti-CCR10 CAR T-cells, we next sought to characterize any potential “on-target, off-tumor” toxicities. Toward this goal, we performed flow cytometry on peripheral blood mononuclear cells (PBMCs) from three independent donors and surprisingly identified that monocytes had noticeable expression of CCR10 (**Fig. 6A**). This finding was unexpected as single-cell transcriptomic datasets indicated no *CCR10* expression on any myeloid cell lineage, including monocytes (**Fig. 1C-D**). Further investigation using the THP-1 cell line as a model for monocytes showed that, despite similar or even higher measured expression of surface CCR10 by flow cytometry than primary monocytes, we saw no cytotoxicity of our CCL27 mutant CAR-Ts, compared to the CD33 positive control CAR-T (**Fig. 6B**).

**Figure 6:**
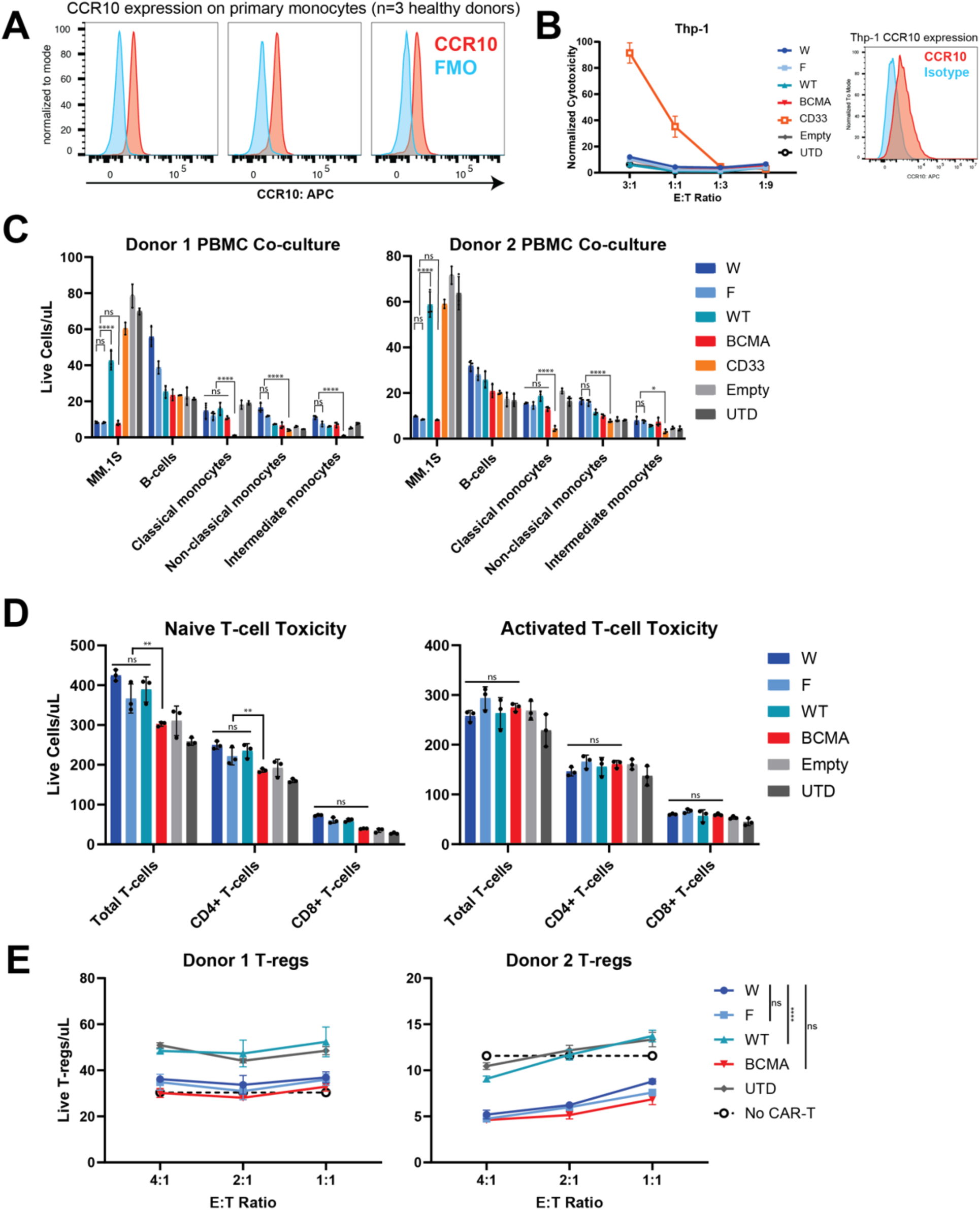
Profiling the toxicity of mutant CCL27-based CAR T-cells. **A.** Flow cytometry of *n* = 3 healthy donor PBMCs focusing on the CD14+ monocytes that show positive staining of CCR10. **B.** Thp-1 monocyte cell line expression of CCR10 (right panel) by flow cytometry, similar to the primary monocytes. *In vitro* cytotoxicity of CCL27 mutant CAR-Ts against Thp-1 monocyte cell line (*left panel*) compared to positive control CD33 CAR-T. **C.** Overnight co-culture of *n* = 2 T-cell donor CAR-Ts with donor-matched healthy PBMCs and MM.1S tumor cells in technical triplicate using flow cytometry with count beads to separate and quantify the mononuclear cell populations (CD19 for B-cells and CD14/CD16 for monocyte subpopulations). CD33 CAR-T is a positive control for monocyte depletion. *p*-values calculated by 2-way ANOVA with Tukey’s correction for multiple comparisons applied: ns = not significant, * = *p*<0.05, & **** = *p*<0.0001. **D.** Overnight co-culture of CCL27 mutant CAR-Ts with donor-matched naïve T-cells or activated T-cells labeled with Cell Trace Violet dye to separate from CAR-Ts, analyzed and quantified by flow cytometry with counting beads separating for CD4+ and CD8+ T-cells. *p*-values calculated by 2-way ANOVA with Tukey’s correction for multiple comparisons applied: ns = not significant & ** = *p*<0.01. **E.** Overnight co-culture of *n* = 2 expanded peripheral T-regs from healthy donors with CCL27 mutant CAR-Ts and MM.1S tumor cells (see Supplementary Data Fig. 5), separated and quantified by flow cytometry with counting beads by labeling CAR-Ts with Cell Trace Violet dye. *p*-values calculated by 2-way ANOVA with Tukey’s correction for multiple comparisons applied: ns = not significant & **** = *p*<0.0001.

Interestingly, CRISPR-based knockout of *CCR10* in the THP-1 cell line, confirmed at the genomic level by the Synthego Interference of CRISPR Edits (ICE) tool^41^, minimally impacted measured CCR10 by flow cytometry; CCL27 mutant CAR T-cells continued to be unable to kill *CCR10* KO THP-1 cells (**Supplementary Data Fig. 5A-C**). Taken together, the combined genomic and functional evidence suggested to us that CCR10 was, in fact, not expressed on monocytes and that the CCR10 antibody (1B5 clone) likely has non-specific binding, leading to an artifactual signal on monocytes. This is known to be a common issue for antibodies against chemokine receptors^42,43^. To assess the more relevant primary PBMC populations, we performed two donor-matched PBMC co-cultures and saw no significant toxicity of the CCL27 mutant CAR-Ts against CD19+ B-cells and CD14+/CD16+ monocyte subpopulations, while robust anti-tumor activity was maintained against the MM.1S tumor cells spiked into the co-culture as a positive control of CAR-T function (**Fig. 6C**). As expected, another positive control CD33 CAR-T strongly depleted classical monocytes (**Fig. 6C**). Surprisingly, we saw no depletion of either activated or naïve T-cells, despite needing to knockout *CCR10* in CAR-T manufacturing to see activity in our previous study^12^, to likely prevent fratricide and/or cross-activation (**Fig. 6D**). To investigate these toxicities in the potentially inflammatory tumor microenvironment, we repeated our co-culture against PBMCs activated with lipopolysaccharide (LPS) and yet again saw no evidence of cytotoxicity of the CCL27 mutant CAR-Ts against either activated B-cells or monocytes (**Supplementary Data Fig. 5D**). Next, we profiled the toxicity against peripheral T-regs isolated and expanded from two healthy donors, finding that, by flow cytometry, they also appear to express CCR10 (**Supplementary Data Fig. 5E**). However, analogous to our data in monocytes, the CCL27 mutant CAR-Ts did not deplete peripheral T-regs while robustly killing MM.1S tumor cells, appearing equivalent to the anti-BCMA CAR-Ts (**Fig. 6E & Supplementary Data Fig. 5F**), and again suggesting an artifact of antibody non-specificity in flow cytometry. Lastly, we performed flow cytometry on CD34+ hematopoietic stem and progenitor cells (HSPCs) from five donors and found no expression of CCR10 (**Supplementary Data Fig. 5G**). Though further characterization of the toxicities of our CCL27 mutant CAR-Ts will be required for clinical translation, these results are encouraging to demonstrate minimal probability of toxicity to critical hematopoietic cells.

## Discussion

Here we present the first, to our knowledge, utilization of a computational structure-guided engineering approach to design a natural ligand CAR-T against any target. We specifically apply our methodology to our recently identified therapeutic target for multiple myeloma, CCR10. We further validated the potential of CCR10 as an immunotherapy target with consistent expression on primary myeloma cells and with potential upregulation in high-risk patient tumors. We then developed enhanced CCL27-based CAR-Ts against CCR10, utilizing computational modeling of the CCL27-CCR10 binding interface to identify two CCL27 mutants that improved the efficacy of our CAR-Ts from almost zero to near the clinically approved cilta-cel anti-BCMA CAR-T.

Comprehensive toxicity testing validated the safety of our CCL27-based CAR-Ts in the peripheral blood compartment.

This work describes a novel potential therapeutic for multiple myeloma patients who relapse after BCMA CAR-T therapy. As we show here, expression of *CCR10* is higher in relapsed myeloma tumors, and higher expression is correlated with worse patient outcomes. Our CCL27-based CAR-Ts against CCR10 may thus be uniquely positioned for this patient population that is most in need. Additionally, this work illuminates the ability to inform CAR-T binder design by integrating computational structural modeling tools with the literature. Most importantly, we show that this approach has the potential to take a very poor-performing CAR T-cell (WT-CCL27) and, with a single round of screening just a handful of variants, identify N-terminal extension mutations (W-and F-CCL27) that significantly improve the efficacy. Importantly, this N-terminal extension could not be identified by the standard antibody engineering strategy of affinity maturation. This natural ligand approach is particularly valuable for chemokine receptors, where the development of antibodies with minimal to no cross-reactivity amongst other homologous chemokine receptors is difficult^42,44^. This limitation was very apparent in our study with the commercially available CCR10 antibody (1B5 clone), which showed non-specific staining of monocytes and potentially peripheral T-regs.

In terms of the limitations of our study, despite the substantial improvement over WT CCL27, our mutant CCL27 CAR-Ts may not have the same durability of response *in vivo* as cilta-cel-based anti-BCMA CAR-Ts. Additionally, these mutant CCL27 CAR-Ts may be less potent against tumors with lower antigen densities of CCR10. In future work, modifications to the intracellular signaling domain^45^ of the CAR and the addition of genetic potency enhancements^46,47^ are planned to attempt to increase the *in vivo* performance of these engineered CCL27-based CAR Ts prior to clinical translation.

In general, the ability to engineer natural ligand-based CAR-Ts beyond physiological constraints using computational structural modeling tools enables targeting traditionally difficult-to-target surface proteins, like chemokine receptors, as done in this work, and other GPCR family members. Beyond CCR10, other chemokine receptors have been identified as therapeutic targets, including CCR4, CCR8, CCR5, CXCR4, and many more^48,49^. Despite the wealth of potential therapeutic targets, the chemokine receptor family has only a few therapeutics approved against it due to the difficulty of identifying binders^50^. Thus, the ability to use this computational structure-guided design strategy on natural ligand binders, illuminated in our work here with CCR10 to other chemokine receptor targets, could have a meaningful therapeutic impact across indications, including various types of cancer, HIV, and inflammatory disorders^49^. Additionally, the unique binding modality of chemokines is particularly intriguing in the context of a CAR T-cell binder because they bind deep into the transmembrane regions of their receptor^51^, enabling a close immunological synapse that may not be achievable with traditional antibody-based binders. The size of the immunological synapse has been shown to significantly impact CAR T-cell efficacy, with a narrow intermembrane space physically excluding the bulky inhibitory protein, CD45^52^.

In conclusion, our work demonstrates that CCR10 is a promising therapeutic candidate for multiple myeloma, and our engineered mutant CCL27 CAR-Ts are an important step in the development of a novel therapy against CCR10 for patients who relapse on BCMA CAR-Ts and need new therapeutic options.

## Funding

This work was supported by funding from the Silicon Valley Community Foundation Myeloma Solutions Fund (to A.P.W. and B.G.B.), UCSF Living Therapeutics Initiative (to A.P.W.), UCSF Stephen and Nancy Grand Multiple Myeloma Translational Initiative (to A.P.W.), and Multiple Myeloma Research Foundation Translational Accelerator Award (to A.P.W.). Salary support was provided by a Multiple Myeloma Research Foundation fellowship (to B.P.E.) and NIH Medical Scientist Training Program grant T32GM141323 (supporting A.K.). Murine studies were performed at the UCSF Helen Diller Family Comprehensive Cancer Center Preclinical Therapeutics Core facility, and flow cytometry/sorting was performed with the UCSF HDFCCC Laboratory for Cell Analysis, both supported by NCI P30CA082103.

## Contributions

N.C., B.P.E., and A.P.W. designed and conceived the study. N.C., B.P.E., E.Y.C., J.M., H.J., and A.K. performed experiments and analyzed data. B.P.E., N.A., and B.G.B. analyzed patient data and datasets. P.P., F.S., Y.Z., and V.S. performed murine studies.

R.D. and T.K. performed structural protein analysis. N.A. and W.J.K. performed flow cytometry on several primary samples. A.C., A.C., A.D.P., T.G.M., and J.L.W. contributed to obtaining primary samples for functional testing. N.C. and A.P.W. wrote the manuscript. All authors provided feedback and approved the manuscript.

## Competing Interests and Disclosures

Patent application filed related to mutant CCL27-based CAR design described here (N.C. and A.P.W.).

