## Supplemental Figures & Methods for "Structure-guided engineering of CCL27 enhances natural ligand CAR T-cells against CCR10 for multiple myeloma"

### Supplementary Data Figure 1

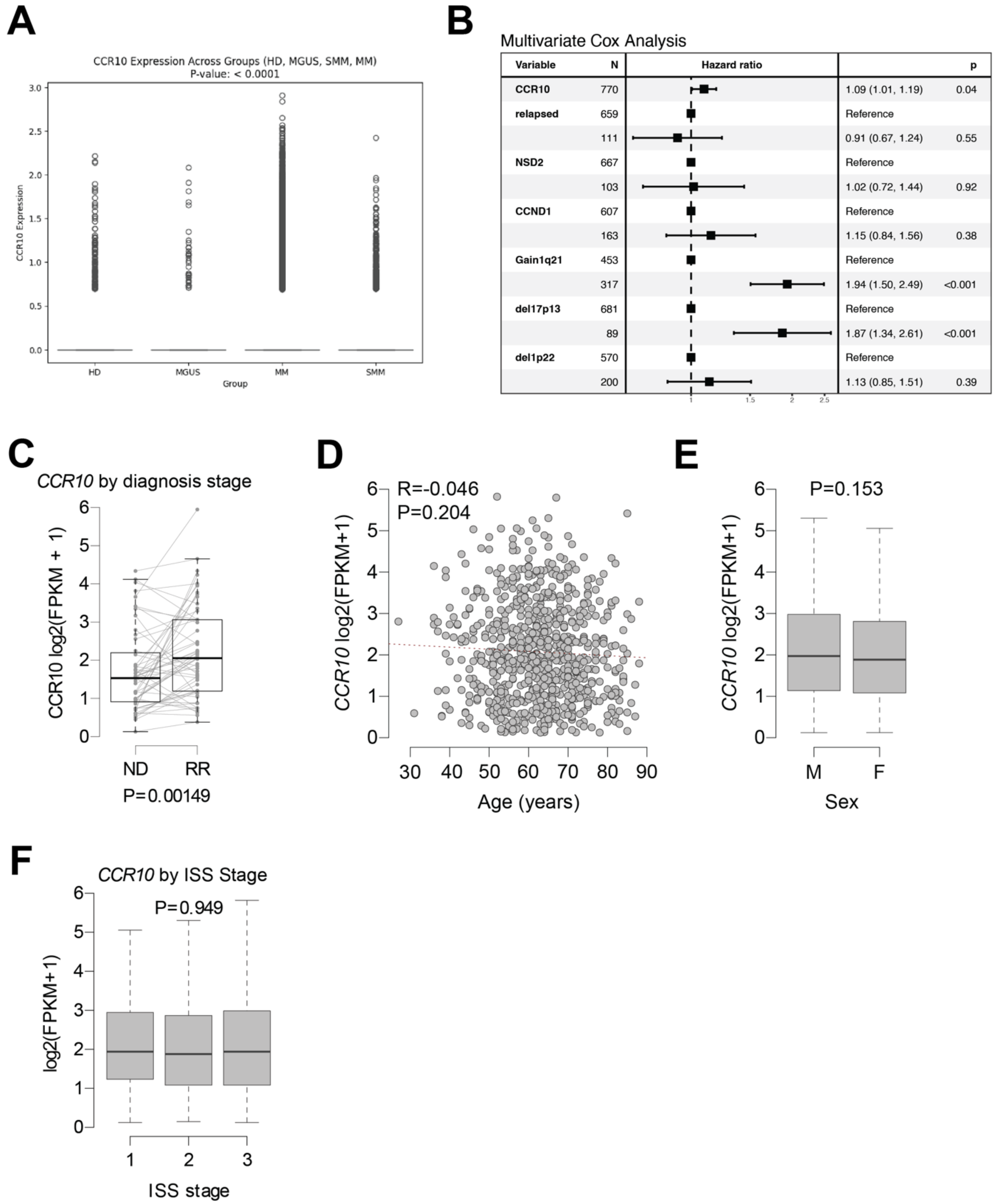


**Supplementary Data Figure 1: Investigation of CCR10 expression in myeloma. A.** Comparison of *CCR10* RNA transcript expression from single-cell RNA sequencing data[1] across myeloma diagnosis stages: “HD” = Health donor, “MGUS” = Monoclonal gammopathy of undetermined significance, “MM” = Multiple myeloma, “SMM” = Smoldering multiple myeloma.

*p*-value calculated by 2-way ANOVA. **B.** Multivariate Cox analysis from the CoMMpass dataset (IA19) to determine the independent risk of *CCR10* expression on myeloma patient outcomes compared to traditional high-risk cytogenetic features. *p*-values calculated by the log-rank test. **C.** Comparison of *CCR10* expression between newly diagnosed (ND) and relapsed/refractory (RR) samples from the CoMMpass dataset (IA21) with matched patient ND and RR samples marked by lines. *p*-value calculated by linear regression with a covariate for patient. **D.** *CCR10* expression by age from the CoMMpass dataset (IA21). *p*-value calculated by linear regression. **E.** *CCR10* expression by gender ("M" = Male and "F" = Female) from the CoMMpass dataset (IA21). *p*-value calculated by linear regression. **F.** *CCR10* expression by ISS (International Staging System) stage for myeloma clinical symptoms severity from the CoMMpass dataset (IA21). *p*-value calculated by analysis of variance with Tukey's post-hoc test.

### Supplementary Data Figure 2

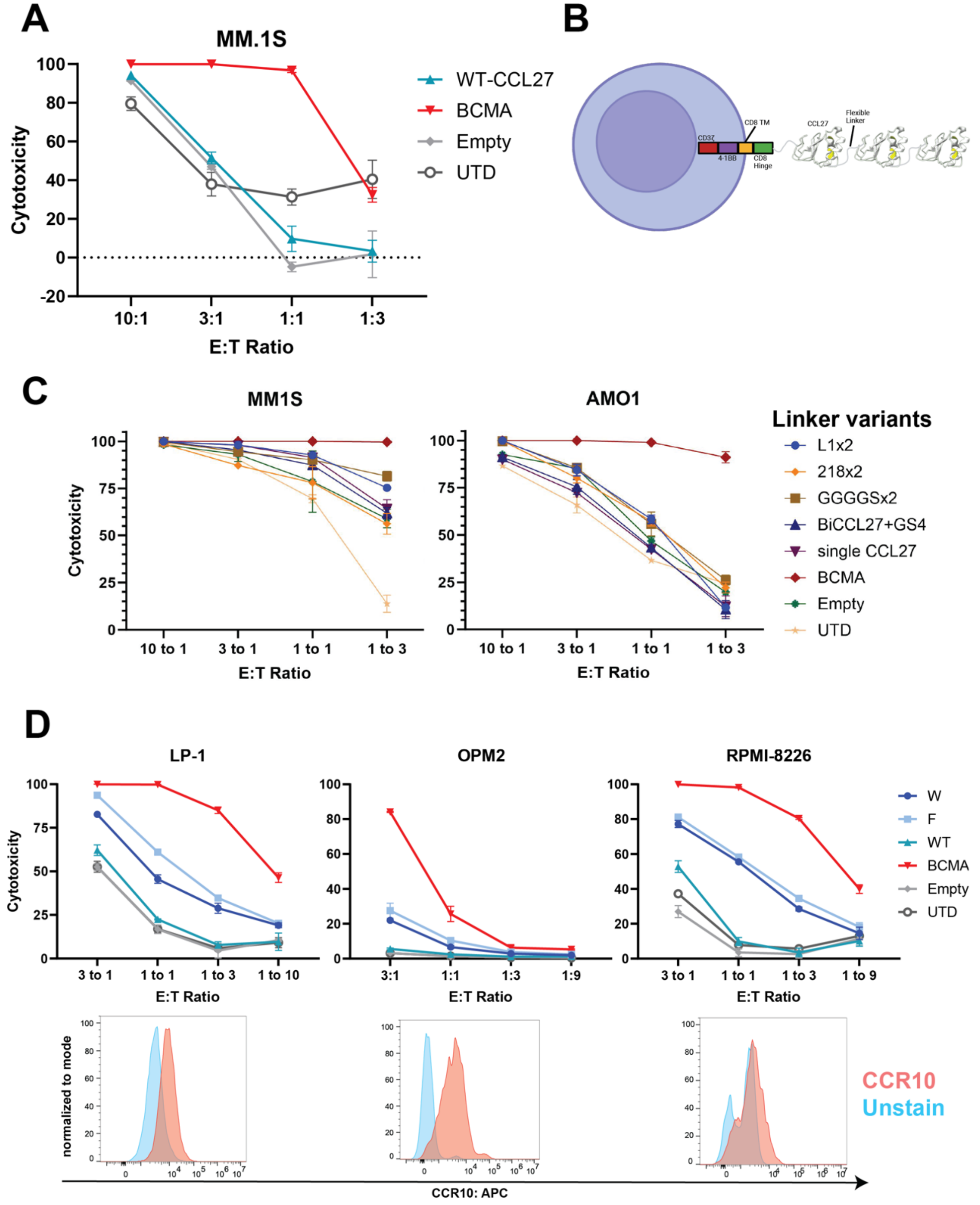


**Supplementary Data Figure 2: Testing various designs of CCL27-based CAR T-cells. A.** *In vitro* cytotoxicity after ~18 hours of co-culture against the MM.1S tumor cell line at the shown E:T (effector to tumor) ratios. *n* = 3 technical triplicates were performed, with the anti-BCMA CAR-T serving as a positive control, while the Empty and UTD CAR-Ts are negative controls. **B.** Schematic of trimeric CCL27 binder connected by flexible linkers that were subsequently screened. CAR-T backbone also shown with CD8 hinge and transmembrane with 4-1BB costimulatory domain and CD3ζ signaling domain. **C.** *In vitro* cytotoxicity of the trimeric CCL27 constructs with different linker variants after ~18 hours of co-culture against the MM.1S and AMO1 myeloma cell lines at the shown effector to tumor (E:T) ratios (*n* = 3 technical replicates) with the same controls as above. **D.** *In vitro* cytotoxicity of the identified CCL27 mutant CAR-Ts after ~18 hours of co-culture with the shown myeloma cell lines (LP-1, OPM2, and RPMI-8226) with the same controls as above (*n* = 3 technical replicates). CCR10 expression measured by flow cytometry is shown in the panels below cytotoxicity plots for the respective cell lines, showing relatively low antigen expression.

### Supplementary Data Figure 3

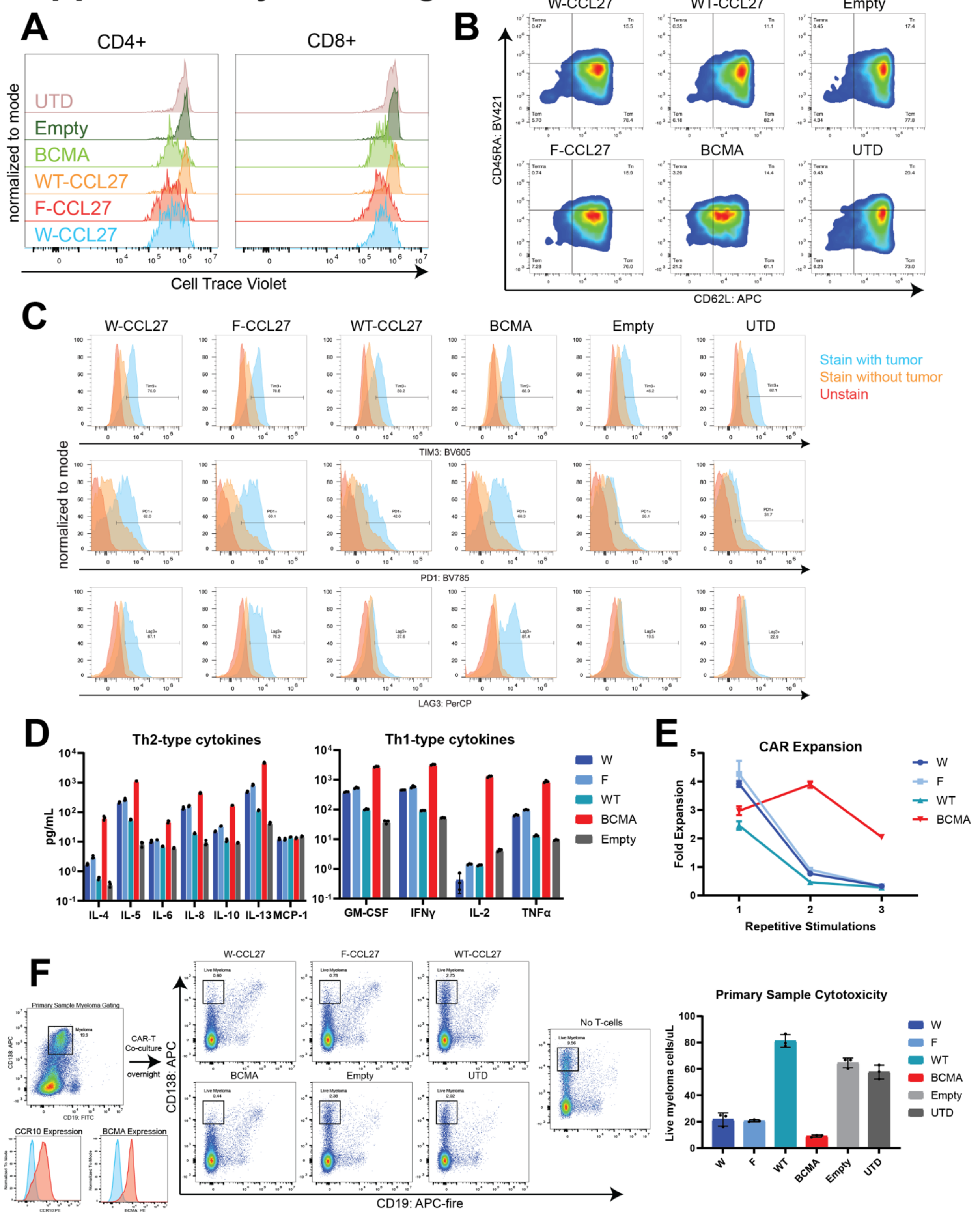


**Supplementary Data Figure 3: Profiling *in vitro* functionality of CCL27 mutant CAR T-cells. A.** Flow cytometry data showing the dilution of the Cell Trace Violet (CTV) dye due to proliferation upon stimulation with MM.1S tumor cells three days before reading out (data shown is for one of three technical replicates; see Fig. 3B for full data). CAR-Ts separated by CD4/CD8 before gating on CTV dye. **B.** Flow cytometry gating of CAR-T Effector/Memory profile 24 hours after antigen stimulation for all six CAR-Ts tested. The data shown is for one of three technical replicates (see main Fig. 3C right panel for full data). **C.** Flow cytometry gating of CAR-T Exhaustion profile 24 hours after antigen stimulation using the markers TIM-3, PD-1, and LAG-3 for all six CAR-Ts tested. Data shown is for one of three technical replicates (see main Fig. 3C left panel for full data). Unstained CAR-Ts are shown in red, stained CAR-Ts without tumor stimulation are shown in orange, and stained CAR-Ts with tumor stimulation are shown in blue. **D.** Concentration of cytokines in the supernatant of a co-culture between CAR-Ts and MM.1S tumor cells at a 1:1 E:T ratio. **E.** Repetitive stimulation of CAR-Ts ($n$ = 3 technical replicates) with tracking of CAR T-cell fold expansion between stimulations by flow cytometry after 48 hours of co-culture (see main Fig. 3D for tumor cell fold expansion). **F.** Representative flow cytometry gating of a myeloma primary sample for tumor burden alongside CCR10 and BCMA expression on the tumor cells (*left panels*). Flow cytometry gating of primary sample/CAR-T co-culture, looking at tumor burden, for one of three technical replicates of the co-culture (*middle panels*). Quantified tumor burden in the primary sample co-culture using counting beads across $n$ = 3 technical replicates at a 4:1 E:T ratio (*right panels*).

### Supplementary Data Figure 4

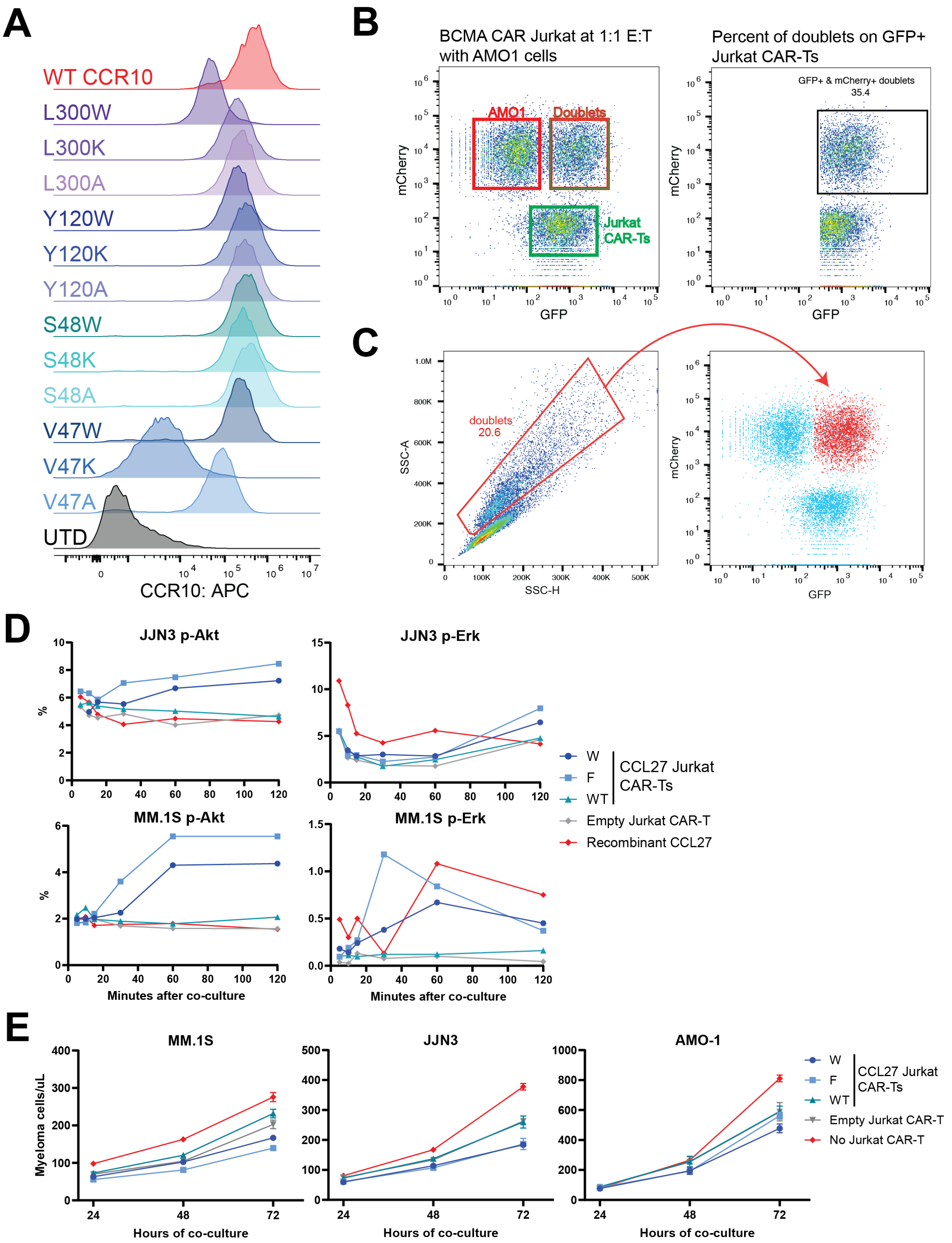


**Supplementary Data Figure 4: Interrogating the mechanism and downstream signaling of mutant CCL27 binding CCR10. A.** CCR10 staining by flow cytometry on the AMO-1 mutant

CCR10 cell lines showing that they all had a similar antigen density (see Fig. 5B for cytotoxicity of CAR-Ts against these cell lines). **B.** Representative flow cytometry gating of doublet assay from the BCMA Jurkat CAR-T at a 1:1 E:T ratio showing clear population of doublets (*left panel*). Representative gating strategy for reporting of percent of doublets on GFP+ Jurkat CAR-Ts that are reported in main Fig. 5D (*right panel*). **C.** Representative flow cytometry plot gating on doublets in the SSC-A vs SSC-H plot (*left panel*). Back gating of doublets from SSC-A vs SSC-H plot onto the mCherry vs GFP plot shows that mainly doublets are mCherry+ & GFP+. **D.** p-Erk and p-Akt flow cytometry staining on the JJN3 and MM.1S myeloma cell lines over the course of 120 minutes. Jurkat CAR-Ts were added to the co-culture at a 3:1 E:T ratio and recombinant CCL27 at 250nM. Values are representative of a single technical replicate. **E.** Co-culture of myeloma cell lines (MM.1S, JJN3, and AMO-1) with Jurkat CAR-Ts at a 3:1 E:T ratio (*n* = 3 technical replicates), with the growth of myeloma cells over 72 hours quantified by flow cytometry using counting beads. No Jurkat CAR-T and empty Jurkat CAR-T conditions were the relevant negative controls for this experiment.

### Supplementary Data Figure 5

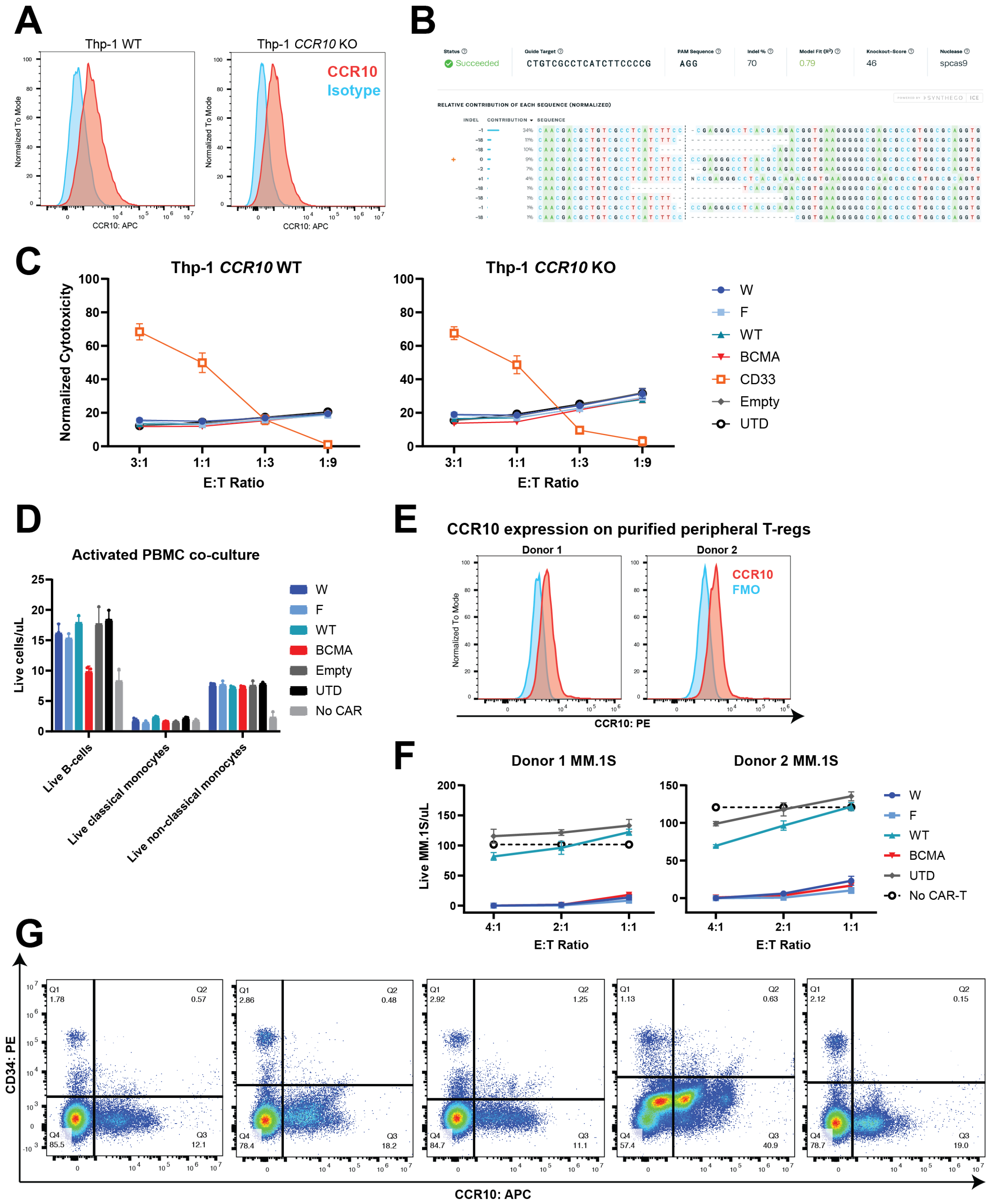


**Supplementary Data Figure 5: Investigation of non-specific CCR10 flow cytometry staining and potential toxicities of CCL27 mutant CAR T-cells. A.** Flow cytometry staining of CCR10 on the THP-1 cell line, with and without *CCR10* KO, showing no meaningful change in

CCR10 staining by 1B5 clone. Representative of two independent knockouts of the THP-1 cell line and flow cytometry experiments on these cells. **B.** Output of Synthego Interference of CRISPR Edits (ICE) tool demonstrates 70% knockout of *CCR10* in the THP-1 cell line based on genomic DNA analysis by Sanger sequencing. **C.** *In vitro* cytotoxicity of CAR-Ts against both THP-1 WT and THP-1 *CCR10* KO cell lines after ~18 hours of co-culture with $n$ = 3 technical replicates at the shown E:T ratios. CD33 CAR-T is a positive control for killing against the THP-1 cell line model of monocytes. **D.** Overnight co-culture of CAR-Ts ($n$ = 3 technical replicates) against PBMCs activated with lipopolysaccharide (LPS), with readout and quantification of relevant PBMC populations using flow cytometry with counting beads. **E.** Expression of CCR10 on purified and expanded peripheral T-regs by flow cytometry ($n$ = 2 donors). See main Fig. 6E for cytotoxicity of CAR-Ts against these T-regs. **F.** Cytotoxicity of CAR-Ts against MM.1S tumor cells ($n$ = 3 technical replicates) in co-culture with T-regs, with readout and quantification by flow cytometry with counting beads. **G.** Flow cytometry on HSPCs from $n$ = 5 GM-CSF mobilized peripheral blood and/or bone marrow samples, showing no CCR10 expression on CD34+ HSPCs.

#### Materials and Methods

##### *List of antibodies*

| Company | Cat # | Lot # | Clone | Antibody | Isotype |
|---|---|---|---|---|---|
| Biolegend | 344814 | B415611 | SK7 | PerCp Human CD3 | Mouse IgG1k |
| Biolegend | 368540 | B419348 | 2D1 | Pacific blue Human CD45 | Mouse IgG1k |
| Biolegend | 363026 | B419907 | SJ25C1 | BV650 - Human CD19 | Mouse IgG1k |
| Biolegend | 367116 | B399433 | 63D3 | FITC Human CD14 | Mouse IgG1k |
| Biolegend | 302012 | B346619 | 3G8 | APC Human CD16 | Mouse IgG1k |
| Biolegend | 369312 | B389996 | 11C3C65 | PerCP Human LAG3 | Mouse IgG1k |
| Biolegend | 345018 | B386958 | F38-2E2 | BV605 Human TIM3 | Mouse IgG1k |
| Biolegend | 329930 | B427545 | EH12.287 | BV785 Human PD1 | Mouse IgG1k |
| Biolegend | 304810 | B411776 | DREG-56 | APC Human CD62L | Mouse IgG1k |
| Biolegend | 384130 | B420339 | HI100 | BV421 Human CD45RA | Mouse IgG1k |
| Biolegend | 352308 | B428363 | DL-101 | APC Human CD138 | Mouse IgG1k |
| Biolegend | 356606 | B371425 | HB-7 | APC Human CD38 | Mouse IgG1k |
| Biolegend | 363030 | B374784 | SJ25C1 | APC Fire - Human CD19 | Mouse IgG1k |
| BD | 567147 | 3160466 | HB7 | FITC Human CD38 | Mouse IgG1k |
| Biolegend | 344840 | B434936 | SK7 | APC Fire Human CD3 | Mouse IgG1k |
| Biolegend | 317416 | B347062 | OKT4 | APC Human CD4 | Mouse IgG1k |
| BD | 564771 | 4012948 | 1B5 | APC Human CCR10 | Mouse IgG2k |
| BD | 550882 | 3128891 | G155-178 | APC Isotype Control | Mouse IgG2k |
| Biolegend | 302008 | B386398 | 3G8 | PE Human CD16 | Mouse IgG1k |
| BD | 561383 | 3110786 | M5E2 | APC Human CD14 | Mouse IgG1k |
| BD | 347313 | 3172576 | SK1 | FITC Human CD8 | Mouse IgG2k |
| BD | 555573 | 2279966 | G155-178 | FITC Isotype control | Mouse IgG2k |
| BD | 552026 | 0286477 | MI15 | PE Human CD138 | Mouse IgG1k |
| BD | 563656 | 4317078 | 1B5 | PE Human CCCR10 | Mouse IgG2k |
| BD | 560378 | 2223482 | M89-61 | PE Human anti pAKT (pS473) | Mouse IgG2k |
| Biolegend | 369510 | B448764 | 6B8B69 | BV421 anti-ERK1/2 Phospho (Thr202/Tyr204) | Mouse IgG2k |
| Biolegend | 343605 | B365415 | 561 | PE Human CD34 | Mouse IgG2k |
| Biolegend | 344722 | B401698 | SK1 | APC Human CD8 | Mouse IgG1k |
| Biolegend | 344620 | B422482 | SK3 | Pacific Blue Human CD4 | Mouse IgG1k |
| Biolegend | 300412 | B341478 | UCHT1 | APC Human CD3 | Mouse IgG1k |
| Biolegend | 357504 | B427225 | 19F2 | PE Human BCMA | Mouse IgG2k |
| BD | 555412 | 4043594 | HIB19 | FITC Human CD19 | Mouse IgG1k |
| Biolegend | 400212 | B367625 | MOPC-173 | PE Isotype control | Mouse IgG2k |
| R&D Systems | FAB193P | AAKZ0520041, AAKZ0623031 | Polyclonal | BCMA PE | |
| BD Biosciences | 557791 | 1109335, 2010314 | SJ25C1 | APC-Cy7 Mouse Anti-Human CD19 | |
| BD Biosciences | 340950 | 2241539, 3094277 | SJ25C1 | CD19 PerCP-Cy5.5 | |
| BD Biosciences | 340673 | 2166696, 3174263 | L27 | CD20 FITC | |
| BD Biosciences | 654664 | 2266689, 2340530 | L128 | CD27 FITC | |
| Cell Signaling | 28034 | 2280897 | E6S8H | CD27 FITC | |
| BD Biosciences | 335808 | 2244920, 2287050, 3068691, 3205822 | HB7 | CD38 PE-Cy7 | |
| Beckman Coulter | A96416 | 200140, 200145, 200146, 200147, 200150, 200154 | J33 | CD45 Krome Orange | |
| BD Biosciences | 340952 | 2032026, 2200861, 3011439 | 2D1 | CD45 PerCP-Cy5.5 | |
| BD Biosciences | 340685 | 2031472, 2166769, 3086543 | MY31 | CD56 PE | |
| Beckman Coulter | A86051 | 230072, 230077, 230078 | 104D2D1 | CD117 APC Alexa 750 | |
| BD Biosciences | 347207 | 2244916, 2280897, 3025565 | MI15 | CD138 APC | |
| BD Biosciences | 564338 | 2056123, 2220654 | 235614 | CD319 APC Alexa 647 | |
| BD Biosciences | 643774 | 2209649, 2265868, 2347785 | TB28-2 | Kappa FITC | |
| BD Biosciences | 642924 | 2223837, 2273943, 2315515 | 1-155-2 | Lambda PE | |

*Flow Cytometry*

Cells were resuspended in FACS buffer (2% FBS in D-PBS) and stained with antibodies for 20-40 min at 4°C, then washed twice and resuspended in FACS buffer. Samples were analyzed using either a Cytoflex Flow Cytometer (Beckman Coulter, Beckman Coulter Navios Flow Cytometer), Attune NXT Flow Cytometer (Thermofisher Scientific), or FACSAria-Fusion (BD Biosciences). Data analysis was completed using FlowJo software, v10.10.0. Compensation was performed with UltraComp eBeads™ Compensation Beads (Invitrogen, 01-2222-42) for antibody fluorophores, Viability Dye Compensation Standard 8μm (Bangs Labs, 451) for amine-reactive viability dyes, and GFP BrightComp eBeads™ Compensation Beads (Invitrogen, A10514) for GFP.

*Cell lines*

Human cell lines were authenticated by short tandem repeat (STR) analysis and routinely tested for mycoplasma contamination. Cells were grown in RPMI 1640 medium supplemented with 20% fetal bovine serum (FBS) and 100 U/mL penicillin-streptomycin. Cell lines used were RPMI-8226, MM.1S, OPM2, Thp-1 (originally obtained from ATCC); AMO-1, LP-1, JJN3 (originally obtained from DSMZ); KMS-11, KMS-26, KHM-11; KMS-34 (originally obtained from JCRB).

*CoMMpass Analyses*

CoMMpass IA21 outcome and clinical data were provided by the Multiple Myeloma Research Foundation (research.themmrf.org). CoMMpass RNA[2] and genetic analysis[3] were performed similarly to analyses previously described. Briefly, summarized RNA counts were downloaded from the Genomic Data Commons, and gene expression subtypes were determined based on published gene expression signatures from Zhan et al.[4] using methods previously described[5]. Structural variants and mutations were determined by Delly[6] and Mutect2[7], respectively.

*Molecular Cloning and DNA Plasmids*

Genes encoding the different anti-CCR10 (CCL27 mutant) and anti-BCMA (cilta-cel) CAR-Ts and mutant CCR10 sequences were synthesized as gene fragments from Twist Biosciences (South San Francisco, CA). DNA fragments were then cloned into a lentiviral expression vector with a Gibson assembly product (NEB, E2611S). DNA sequencing was performed to confirm the precise integration of the sequences into the backbone. This final construct was then expressed in Stbl3 Competent E. coli (Macro Lab, UC Berkeley, CA). DNA was isolated using either QIAGEN Plasmid Plus Midi Kit (Qiagen, 12943)

*Lentiviral Vector Production*

Lenti-X 293T cells (Takara, Cat # 632180) were transfected with each CAR expression plasmid and Mirus Bio™ TransIT™-Lenti Transfection Reagent (Cat # MIR6600). Lenti-X 293T cells were cultured for 2-3 days, and then lentiviral supernatant was harvested and concentrated using Lenti-X Concentrator (Takara Bio, 631232). Concentrated pellets were resuspended in cold serum-free Opti-Mem media and either stored at -80°C or applied to cells immediately.

*CAR T-cell Production and Expansion*

Primary human T-cells were purified from either leukoreduction filter products of anonymous healthy blood donors from Vitalant (San Francisco, CA) under an institutional reviewer board-exempt protocol in accordance with the U.S. Common Rule (Category 4) or de-identified donor leukapheresis products obtained from StemCell Technologies. CD3+ T-cells were isolated by negative selection using the EasySep™ Human T-Cell Isolation Kit from StemCell Technologies (Catalog # 17951). CD3+ T-cells were thawed, cultured separately in media overnight, and then cells were counted the following day. T cells were cultured in CTS™ OpTmizer medium with CTS supplement (Thermo Scientific, A1048501) supplemented with 5% human AB serum (HP1022; Valley Medical), GlutaMAX (Gibco™. Cat # 35050061), and 100 U/mL penicillin/streptomycin (Fisher Scientific, 15-140-122) and were passaged every 2 days. For expansion, T cells were stimulated with CD3/CD28 Dynabeads (11131-D; Thermo Fisher Scientific) according to the manufacturer's instructions (20 µL of beads per 1 million T-cells) for 5 days and grown in the presence of recombinant interleukin-7 (PeproTech, 200-07) and interleukin-15 (PeproTech, 200-15) at 10 ng/mL. Transduction with CAR lentivirus was performed 24 hours after the start of bead stimulation. After the removal of CD3/CD28 activation beads, transduction efficiency was assessed by flow cytometry. CAR-T cells were labeled with intracellular GFP, so the percentage of GFP+ cells determined the percentage of CAR-T cells generated.

*PBMC Co-culture*

Matched PBMCs were purified from the same de-identified donor leukapheresis products used for CAR-T manufacturing, obtained from StemCell Technologies, using Lymphoprep density gradient separation (StemCell Technologies, 18061) and frozen for storage. CAR-Ts were co-incubated in technical triplicate with thawed donor-matched PBMCs at a 1:1 E:T ratio and MM.1S tumor cells at a 2:1 E:T ratio overnight. Flow cytometry was performed the next day using CD19 to identify B-cells, CD14 to identify monocytes, and CD45/SSC-A to identify MM.1S tumor cells from the co-culture populations. Live and dead cells were separated using the Zombie r718 dye (Biolegend, 423116). Cell counts were quantified and normalized using counting beads (Biolegend, 424902) to get the reported live cells/uL values.

*Murine Studies*

All murine studies were conducted under UCSF Institutional Animal Care and Use Committee-approved protocols. NSG (NOD.Cg-*Prkdcscid Il2rgtm1Wjl*/SzJ, Jackson Laboratories) mice, 6-9 weeks old, were injected intravenously via tail vein with 1e6 multiple MM tumor cells stably expressing luciferase. On day 7 or 8 after tumor injection, mice were injected intravenously with 3e6 of CAR+ T-cells. Bioluminescence imaging was done weekly to assess tumor burden (Perkin Elmer In Vivo Imaging System, Caliper Life Sciences). The survival endpoints of the studies were determined by signs of symptomatic illness in the animals and required veterinary protocols for humane euthanasia.

*In vitro cytotoxicity assays*

All cell lines were engineered to stably express luciferase using lentiviral transduction. For each cell line, 50,000 MM cells were seeded in 50 uL of RPMI20 into a white 96 well plate (Greiner Bio-One). Subsequently, CAR-T cells were added at four different effector:target ratios (3:1, 1:1, 1:3, and 1:9) in 50uL of T-cell media with *n*=3 replicates per ratio and CAR-T construct. This coculture was incubated at 37C for 18-24 hours. The next day, 100uL of D-Luciferin (Gold Biotechnology, LUCK-1G) was added to each well to a final concentration of 375 µg/ml, followed by luminescence detection using GloMax Explorer (Promega). The bioluminescence readings were averaged amongst the *n*=3 technical replicates for each construct and ratio. Bioluminescence readings were logarithmically scaled and normalized to the maximum and minimum values for each cell line to create a cytotoxicity scale of 0-100. Assays were performed in biological replicates with T-cells from at least 2 or 3 donors.

*CAR-T In vitro proliferation*

CellTrace$^{TM}$ Violet (Invitrogen, C34557) was diluted to 5µm in warm PBS (37°C). CAR T-cells were washed in PBS to remove all serum from culture media and resuspended in the CellTrace$^{TM}$ Violet (CTV) staining solution. CAR-Ts were incubated for 20 minutes at 37°C and washed in OpTmizer T-cell media supplemented with human serum to neutralize all unbound dye. Labeled CAR-Ts were incubated with MM.1S tumor cells at a 1:1 E:T ratio in a U-bottom 96-well plate (Greiner Bio-One). Flow Cytometry was performed 72 hours after the initial co-culture was set up, gating on CD3+ and GFP+ CAR-Ts, separated by CD4/CD8. Proliferation was characterized as “% Divided Cells”, gated as the dilution of the CTV dye as compared to CAR-Ts that did not receive tumor stimulation.

*CAR-T cell in vitro immunophenotyping*

CAR-T cells were cultured with tumor for 24 hours at a 1:1 E:T ratio followed by flow cytometry profiling for exhaustion and memory markers. The markers used were CD3, CD45RA, CD62L, TIM3, LAG3, and PD1 (see table of antibodies for clone information). Data was analyzed and processed using FlowJo 10.10.0. Memory phenotype of CAR-T cells was determined using the following characteristics: stem cell memory ($CD62L^+CD45RA^+$), central memory ($CD62L^+CD45RA^-$), effector memory ($CD62L^-$ $CD45RA^-$), and effector T cells ($CD62L^-$ $CD45RA^+$). Exhaustion was characterized by the percent of cells positive for each individual exhaustion marker and also the frequency of triple-positive cells.

*Structural Modeling with AlphaFold3 and Rosetta*

AlphaFold3 was utilized to model the binding interaction of CCL27 and any of its mutants with CCR10. These predicted structural models were visualized and analyzed with UCSF ChimeraX[8], developed by the Resource for Biocomputing, Visualization, and Informatics at the University of California, San Francisco, with support from National Institutes of Health R01-GM129325 and the Office of Cyber Infrastructure and Computational Biology, National Institute of Allergy and Infectious Diseases. Rosetta analysis was performed with the Flex ddG protocol[9], starting with the AlphaFold3 prediction of the G-CCL27 structure and making subsequent mutations to the N-terminal amino acid extension. The Rosetta fa_talaris2014 score function was utilized to calculate the reported ΔΔG (total score) and total van der Waals energy (fa_atr + fa_rep) of the mutations to the N-terminal amino acid extension of CCL27 across the several structures generated by the Flex ddG protocol.

*Supernatant Cytokine Analysis*

CAR-Ts were cocultured with the MM.1S tumor cell line in technical triplicate at 1:1 E:T ratio. The supernatant was collected, diluted 1:1 with RPMI media and snap frozen using liquid nitrogen. The analysis of cytokine levels was performed by Eve Technologies Corp. (Calgary, Alberta), using Luminex xMAP technology for multiplexed quantification. Multiplexing analysis was done with the Luminex™ 200 system (Luminex, Austin, TX, USA). Fifteen markers were simultaneously measured in the samples using Eve Technologies' Human Focused 15-Plex Discovery Assay® (MilliporeSigma, Burlington, Massachusetts, USA) according to the manufacturer's protocol.  The markers tested included: GM-CSF, IFNγ, IL-1β, IL-1RA, IL-2, IL-4, IL-5, IL-6, IL-8, IL-10, IL-12p70, IL-13, IL-17A, IL-23, TNF-α. Each marker's sensitivity values are available in the MilliporeSigma MILLIPLEX® MAP protocol.

*Repetitive Stimulation Assay*

CAR T-cells were co-cultured with MM.1S tumor cells at a 1:1 E:T ratio and left for 48 hours before assaying by flow cytometry using counting beads (Biolegend, 424902) to determine the absolute counts of CAR-T and tumor cells in the co-culture. This data was used to calculate the number of MM.1S tumor cells necessary to add to the co-culture to reestablish a 1:1 E:T ratio. This protocol was repeated for three cycles of stimulation, with the flow cytometry data being used to track CAR-T and tumor cell expansion between stimulations.

*Primary Sample Cytotoxicity Assays*

Primary samples were collected fresh under University of California, San Francisco (UCSF) Institutional Review Board-approved protocols and in accordance with the Declaration of Helsinki. Bone marrow samples were processed by red blood cell lysis and promptly analyzed by flow cytometry for CD138+/CD19- myeloma cells and expression of CCR10 and BCMA. Samples with appreciable tumor burden (15%+) and positive for CCR10 and BCMA were set up in overnight co-cultures with CAR T-cells. The following day, co-cultures were analyzed by flow cytometry for tumor burden (CD138+/CD19- myeloma cells) and quantified with counting beads (Biolegend, 424902).

*CRISPR-Cas9 Gene Knockouts*

Cas9 protein and target or scramble sgRNA were mixed in a 1:2.5 molar ratio (Synthego Corporation) and incubated at 37 °C for 10–15 min. 1e6 of either primary CD3+ T cells or MM cell lines were spun down and washed with PBS, resuspended in 20 μL of P3 nucleofection/solution plus cas9/sgRNA mixture (P3 Primary Cell 4D-NucleofectorTM X Kit S) for primary T cells or SF Cell Line 4D-NucleofectorTM X Kit S (Lonza) for MM cell lines, and nucleofected using EO-115 or DS-137 nucleofection program in Lonza 4D-Nucleofector, respectively. Immediately after nucleofection, 80 μL of warm (CTS™ OpTmizer™ T Cell Expansion SFM, Gibco™) or RPMI 1640 media (Gibco) supplemented with 20% FBS (RPMI20) was plated into each well and incubated at 37 °C for 15 min, then transferred to media supplemented with IL-7 and IL-15 for T-CD3+ T cells and RPMI20 media for cell lines to recover. Then, based on the surface expression by flow cytometry, negative clones were sorted using FACSARIA-Fusion or FACSARIA III flow cytometer (BD Biosciences). The sgRNA sequence that was obtained from the Brunello Library is as follows: CCR10: CUGUCGCCUCAUCUUCCCCG. Scrambled sgRNA specified by Synthego Corporation.

*Doublet Assay*

Jurkat T-cells were engineered to express the CAR-T binders using the same lentiviral constructs used to make CAR-Ts from primary T-cells to make Jurkat CAR-Ts. The doublet assay protocol was followed similarly to the initial paper[10], with a brief summary of methods following. Myeloma cell lines and Jurkat CAR-Ts were resuspended in fresh RPMI20 and seeded in a cell culture-treated 96-well U-bottom plate (Falcon, 353077) at the indicated E:T ratios, ensuring that the total number of cells in the well was at 50,000 for all ratios. Cells were co-cultured at 37°C for 30-60 minutes to allow binding/doublet formation and immediately run on the flow cytometer with separation of Jurkat CAR-Ts (GFP+) from myeloma cell lines (mCherry+) and binding interactions being GFP+ & mCherry+.

*Calcium flux*

Myeloma cell lines were washed in PBS to remove all serum from the culture, and Fura Red™ (Invitrogen, F3021) calcium flux dye was diluted in PBS warmed to 37°C to 2µM to make the dye staining solution. Myeloma cells were subsequently incubated with Fura Red dye staining solution at 37°C for ~30 minutes. After incubation, cells were washed in RPMI20 media to neutralize all free-floating dye. Stimulants (3:1 E:T Jurkat CAR-Ts, 500nM Recombinant WT-CCL27 working concentration, and 5ug/mL Ionomycin working concentration) were prepared. Of note, Jurkat CAR-Ts were utilized to better represent the number of CCL27-CCR10 interactions as compared to recombinant CCL27. Dye-labeled myeloma cells were run on the flow cytometer for ~30 seconds to establish a baseline calcium signal, and then the stimulant was subsequently added and immediately measured for 5 minutes with an event rate at ~4,000 events per second. Jurkat CAR-Ts (GFP+) were separated from the dye labeled myeloma cell lines (mCherry+) during analysis. Ratiometric analysis of the Fura Red calcium flux dye[11] signal of labeled myeloma cell lines (MM.1S and JJN3) was calculated as the signal off the blue laser ($CA^{2+}$ free) divided by the violet laser ($CA^{2+}$ bound). Values reported depict the moving average of the Fura Red ratio over time (~5 min) of a single replicate but are representative of three independent experiments.

*p-Erk and p-Akt staining*

Myeloma cell lines (MM.1S and JJN3) were prepared in fresh RPMI20 and seeded in a U-bottom 96-well plate. Jurkat CAR-Ts and recombinant CCL27 were prepared at twice their working concentration (3:1 E:T and 250nM respectively) in RPMI20 and added to the myeloma cell lines at the indicated time points, such that all cells were fixed at the same time. Cells were fixed with fixation buffer (Biolegend, 420801) that was warmed to 37°C. Cells were subsequently permeabilized using True-Phos™ Perm Buffer (Biolegend, 425401) chilled to -20°C. Subsequent processing was performed following the manufacturer's protocol for True-Phos™ Perm Buffer. Co-cultures were run on the flow cytometer with separation of Jurkat CAR-Ts (GFP+) and the target myeloma cells (mCherry+).

*scRNAseq analysis for CCR10*

Single-cell RNA data analysis from GSE223060 and previously published[12] was conducted in R (v4.1.2) with Seurat (v4.3.0)[13]. Samples with more than 25% of plasma cells were selected for the analysis. (MMRF_1325, MMRF1537, MMRF1640, MMRF_2038, MMRF_1267, MMRF_1720, MMRF_1505, MMRF_2251, MMRF_2259). Quality control excluded cells expressing fewer than 500 genes, fewer than 1,000 total RNA counts, more than 50,000 features, or with >10% mitochondrial content. Data were normalized, and highly variable features were identified for downstream analysis. Batch effects were corrected with the Harmony[14] package in RStudio (v0.1.1), grouping by experimental condition, and embeddings were used for UMAP visualization and clustering (resolution = 0.5). Gene symbols were annotated using biomaRt. Differentially expressed genes (DEGs) between clusters were identified with a log fold change threshold >0.25 and filtered for average log fold change >0.5. Visualization was performed with ggplot2[15] (v3.5.1) and gridExtra (v2.3)[16].

For the analysis of monocytes, we obtained a publicly available single-cell RNA-sequencing dataset from a collection of scRNA-seq and scTCR-seq data from more than 200 samples from 182 patients with multiple myeloma, MGUS, and SMM, and non-cancer controls[1] (https://zenodo.org/records/13646014). Data was primarily collected from Gene Expression Omnibus (GEO) Maura et al. under accession GSE161195[17,18] Bailur et al. (GSE163278[19]).Oetjen et al. (GSE120221[20]). Granja et al. (GSE139369[21]), Zavidij et al. (GSE124310[22]), Kfoury et al. (GSE143791[23]), and Zheng et al. (GSE156728[24]). Using a pan-immune collection (panImmune.h5ad), it was analyzed in Python using Scanpy (v1.9) through Google Colab. We subsetted the analysis to bone-marrow myeloid cells and applied quality-control filters, excluding cells expressing fewer than 500 genes, fewer than 1,000 total RNA counts, more than 10,000 features, and mitochondrial content < 10%. Following QC, using default Scanpy normalization, log1p transformation, and PCA, then visualized CCR10 expression by extracting single-cell CCR10 values and plotted group-wise expression distributions with Seaborn boxplots. Analysis code for both Seurat and Scanpy analysis are available at the Github repository [https://github.com/BonellPatinoE/CCR10-in-Myeloma.git].

*Multivariate Cox Proportional Hazards Modeling for CCR10*

Differential gene expression analysis based on cytogenetic risk was performed on RNAseq data from primary MM samples from the Multiple Myeloma Research Foundation (MMRF) CoMMpass study (release IA19), which are available to registered users through the MMRF Researcher Gateway (registration information available at https://mmrf.formstack.com/forms/research_gateway_ registration). The RNAseq data files were loaded and processed through DESeq2 (v1.46.0)[25] in R (v2025.05.0+496), 'apeglm' for LFC shrinkage[26] and "gprofiler2" (v0.2.3) for gene list functional enrichment analysis and namespace conversion[27]. Sample-level cytogenetic metadata, including translocations (NSD2, CCND1) and copy-number alterations (del17p13, gain1q21, and del1p22), were extracted from the provided SeqFISH files and merged by sample identifier. Only bone-marrow ("BM") specimens were retained. Samples missing any metadata or with non-intersecting identifiers were excluded. Using clinical and survival information in CoMMpass, we fit a Cox model using the survival package[28] (v3.8-3) for cytogenetic variables, and relapse status was coded as factors (levels 0/1). Hazard ratios (HRs) and 95% confidence intervals were extracted from the model summary. Statistical significance was assessed at $\alpha = 0.05$, two-sided. We visualized the multivariate HRs using the forestmodel package[29] (v0.6.2), and composed with ggplot2 (v3.5.1) with all covariates displayed on a single forest plot[30]. The Cox model was fit by maximizing the partial likelihood (with Efron's method for ties), and inference was provided via the Wald test for individual coefficients and the likelihood-ratio and score (log-rank) tests for the overall model fit. Analysis code is available in a repository at [https://github.com/BonellPatinoE/CCR10-in-Myeloma.git].

*Statistics and reproducibility*

All statistical analyses were performed using GraphPad Prism v.9 unless stated otherwise. The data are represented as mean ± S.D., and statistical significance defined as p values of <0.05. Applied statistical tests are noted in the relevant figure legend. All n values given are biological replicates, unless otherwise specified. Data distributions are assumed to be normal, but not formally tested. No data were excluded from the analyses. Mice were randomized based on body weight before treatment. The Preclinical Core Facility staff was blinded to mouse treatment arms and relevant outcomes. The other investigators were not blinded to allocation during the experiments and outcome assessment.

*Data Availability*

Code for CoMMpass analysis and scRNA-seq analyses available at:
https://github.com/BonellPatinoE/CCR10-in-Myeloma.git

30. ggplot2: Elegant Graphics for Data Analysis | SpringerLink. https://link.springer.com/book/10.1007/978-0-387-98141-3.
